# Mosaic terrestrial diversity dynamics through the Permo-Triassic interval

**DOI:** 10.64898/2026.04.09.717602

**Authors:** Bingcai Liu, Kai Wang, Yao Wang, Honghe Xu

## Abstract

The end-Permian mass extinction (EPME) represents the most severe biotic crisis of the Phanerozoic Eon on Earth and has been well documented in marine taxa. However, its impact on terrestrial organisms and ecosystems remains incompletely understood. Here we present a high-resolution reconstruction of terrestrial diversification dynamics and spatial reorganization across the Permo-Triassic boundary (PTB) using comprehensive occurrence data of macroplants, sporomorphs and vertebrates. Terrestrial responses to the EPME show highly temporal, regional and taxonomic heterogeneities. Plants experienced a genus-level diversity loss of ∼ 6.7%, across the PTB, whilst vertebrates, a lagged decline from the late Permian, peaking at a diversity loss of ∼ 66.7%. Global distributions of plant and vertebrate show converging on similar latitudinal gradients post the PTB. Plant diversity loss is disproportionately high in low-latitude and tropical regions and progressively lower toward mid- and high-latitudes. Our study facilitates a fine-grained understanding to terrestrial macroevolution in geologic history through multi-analysis of a large volume of fossil data. Our findings challenge the long-held notion of global terrestrial collapse and mass extinction in plants during the PTB and offer a deep-time analogue for uneven response of extant terrestrial biodiversity to ongoing climate change.

## INTRODUCTION

As the most severe biodiversity crisis in the Phanerozoic, the EPME wiped out ∼ 81% or 95% species of marine organisms within a short time bin of 60 ± 48 kyr (Sepkoski, 1978; Alroy et al., 2008; Alroy, 2010; Burgess et al., 2014; Stanley, 2016). Such dramatic biodiversity loss was recognized based on abundant marine fossil taxa from continuous strata with fine dating and correlation through the PTB. However, controversies have been raised on 1) asynchronous geochemical data with the EPME event (Fielding et al., 2019; Zhang et al., 2016; 2021; Wu et al., 2024; He et al., 2024; Chu et al., 2025), and 2) whether terrestrial organisms underwent the genuine great extinction (Chu et al., 2025; Benton, 1995; Cascales-Miñana et al., 2016; Lucas, 2017; Mays et al., 2021; Viglietti et al., 2021) or no mass extinction at all (Niklas et al., 1983; Niklas et al., 1985; Nowak et al., 2019; Paterson et al., 2024; Peng et al., 2025; Schneebeli-Hermann and Galasso, 2025).

Precise depiction of terrestrial diversity dynamics through the PTB is necessary to understand the EPME and requires a variety of technical means. Relatively, fossil records of terrestrial plants and vertebrates are readily influenced by regional variability and depositional/preservation biases, and are more complicated to interpret. For example, plant diversity inferred from different areas, such as Angara, Euramerica, South China, and Gondwana, appear inconsistent trajectories and patterns, reflecting ecological, regional and depositional variations in plant fossil records (Rees, 2002; Fielding et al., 2019; Nowak et al., 2022; Peng et al., 2025).

Previously, regional- and global- scaled datasets have been assembled to investigate terrestrial organisms’ responses to the EPME, advancing our understanding of terrestrial ecosystem changes during this interval (Looy et al., 1999; Rees et al., 2002; Cascales- Miñana et al., 2016; Lucas et al., 2017; Benton et al., 2018; Fielding et al., 2019; Nowak et al., 2019; Wu et al., 2024; Peng et al., 2025). However, difference in analytical approaches and limitation in data collection hindered detailed portraying terrestrial diversity dynamics across the PTB. New insights into terrestrial diversification during the EPME are potentially to uncover utilizing a robust statistical framework to address a series of issues specific to terrestrial fossils and strata, such as inaccurate correlation, preservation biases, and data incompleteness (Fielding et al., 2019; Wu et al., 2024; Nowak et al., 2019; Peng et al., 2025; Sues and Fraser, 2010). Here, we overcome this challenge and facilitate to precisely depict terrestrial ecosystem responses to the EPME by compiling a global dataset comprising occurrence records of 12,936 macroplants, 52,486 sporomorphs and 1,609 terrestrial vertebrates across the PTB and comprehensive analyses taking into account of low-resolution terrestrial stratum, dispersal of land organism, biotic and abiotic factors affecting diversity, latitudinal gradient of distribution, and biogeographic zonation. We reveal the heterogeneous diversity dynamic varying in temporal, regional and taxonomic terrestrial organisms through the PTB, namely the mosaic terrestrial diversification pattern. A series of biotic and abiotic factors underlying the pattern are comprehensively assessed.

## RESULTS

### Diversity of terrestrial organisms through the PTB

Macroplant exhibits increases in both origination (mean *λ*=0.2614, from 253-251 Ma) and extinction (mean *µ*=0.2424, 254-251 Ma) rates from the late Wuchiapingian to Induan (Figure 1A), during which the net diversification rate maintains transient declined (Figure 1B). The genus-level diversity decreased by 5.7% from late Wuchiapingian to early Changhsingian (255-253 Ma) (Figure 1C), followed by an increase during late Changhsingian. Diversity then decreases by a further 6.7% from the PTB to Olenekian (251.4-249 Ma) and remains relatively high and stable through late Olenekian to Anisian (Figure 1C).

**Figure 1.**
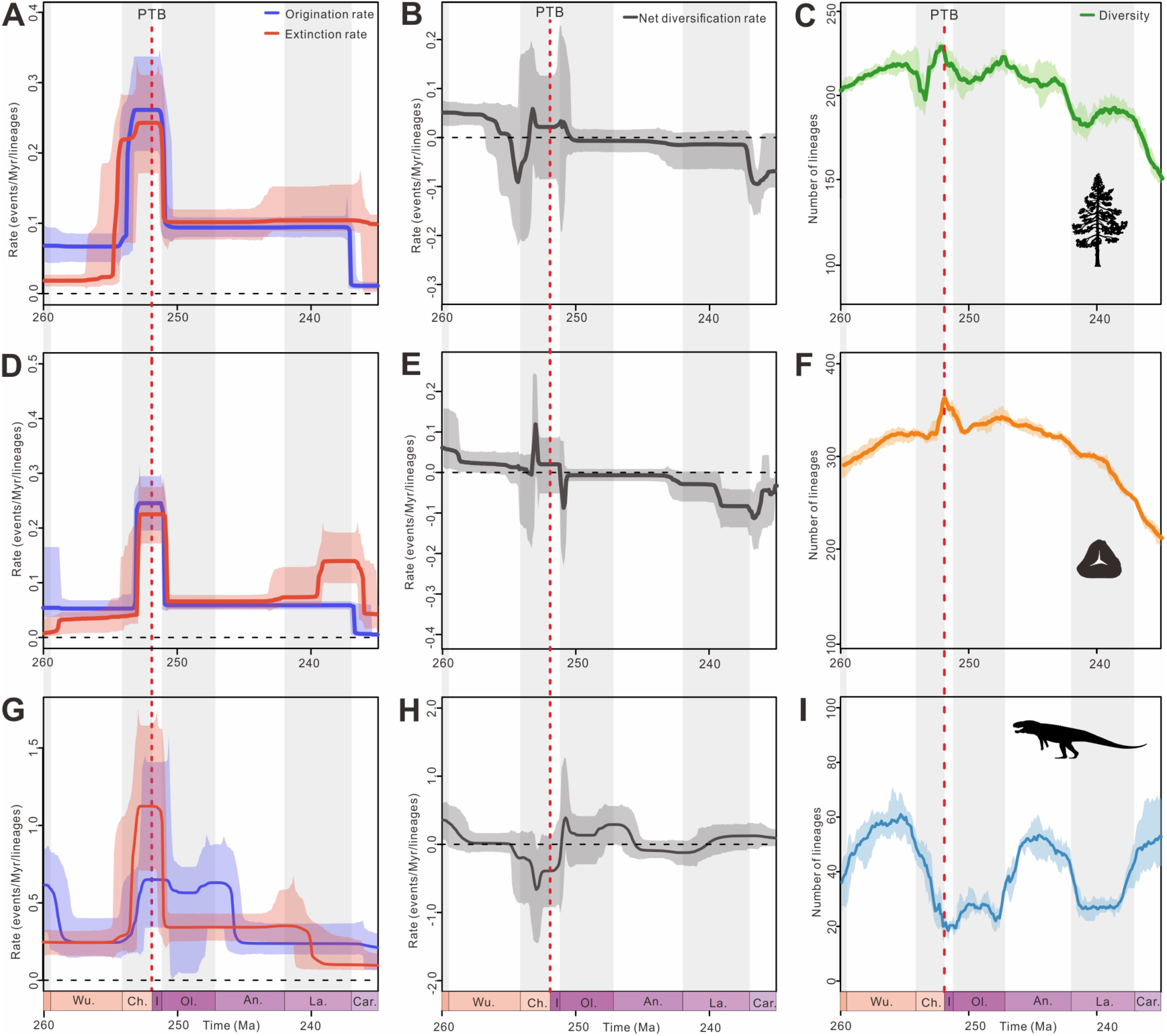
Estimated diversification and diversity dynamics of major groups of terrestrial ecosystems in genus level. Origination rate, extinction rate, diversification rate and diversity of macroplants (A-C), sporomorphs (D-F), terrestrial vertebrates (G-I) through the Permian Triassic Boundary (PTB). Solid lines indicate mean posterior density and the shaded areas show 95% highest posterior density (HPD) intervals. Wo. Wordian, Cap. Capitanian, Wu. Wuchiapingian, Ch. Changhsingian, I. Induan, Ol. Olenekian, An. Anisian, La. Ladinian, Car. Carnian. The silhouettes of plant and vertebrate were taken from https://www.phylopic.org/. The diversification and diversity dynamics of macroplants and vertebrates in species level see Figure S2.

Major plant clades exhibit differentiated diversity patterns. Coniferophyta, Cycadophyta, Ginkgophyta and Lycophyta maintain stable diversity across the PTB (Figure S3A-F). The diversity of Cycadophyta and Pteridophyta continues to increase from Wuchiapingian to Olenekian (Figure S3B, F). Only groups of Pteridospermatophyta and Sphenophyta show high declines of 22.2% and 40.0%, respectively from Changhsingian to Olenekian ages (Figure S3G, H). Such levels of diversity loss do not qualify as a mass extinction (Sepkoski et al., 1981; Hull et al., 2015).

Sporomorphs show increasing origination (mean *λ*=0.2453, 253-251 Ma) and extinction rates (mean *µ*=0.2250, from 252-250 Ma) during late Wuchiapingian to Induan (Figure 1D), high net diversification rate in early Changhsingian (Figure 1E) and a short rise in diversity in late Changhsingian (Figure 1F). Then, their net diversification rate experiences a temporary negative shift after the PTB, and their diversity declines 8.6 % in the earliest Triassic (from 251.6-249.8 Ma), but recovers soon afterward. Diversity of both macroplant and sporomorph show similar pattern and no major collapse is seen in genus level (Figure 1A-F). Whist at the species level, macroplant displays moderate fluctuations in origination and extinction rates, with a short negative excursion and 19.8 % species loss at the PTB (252-249 Ma) (Figure S2A-C).

Terrestrial vertebrates experience a pronounced evolutionary turnover. At the genus level, their origination (mean *λ*=0.6433, from 252-250 Ma) and extinction (mean *µ*=1.1022, from 253-251 Ma) rates both increase sharply, with extinction rates markedly exceeding origination rates (Figure 1G), leading to a substantial negative shift of net diversification rates (−0.6626) and a significant reduction ∼66.7 % in generic diversity from the late Wuchiapingian to late Changhsingian (257-251 Ma) (Figure 1H and J). At the species level, the diversification of vertebrates shows the similar pattern to the genus level (Figure S2D-F).

### Spatial variation and latitudinal trajectory in terrestrial ecosystems

The spatial variations of the terrestrial ecosystems are reflected by the spatio-temporal distribution of macroplants and vertebrates from late Permian to Middle Triassic. Macroplants are widely distributed in a range of climatic zones, spanning cool and warm temperatures, arid, cool and tropical belts (Boucot et al., 2013). Large number of macroplants are concentrated in tropical and warm temperature climatic zones at low latitudes in the Lopingian paleogeographic map (Figure 2E). Following the PTB, the distribution of macroplants contracts markedly during Induan and Olenekian ages and shranks sharply in tropical regions as the tropical climatic zone contracted, especially in the South China Paleoblock at low latitudes (Figure 2).

**Figure 2.**
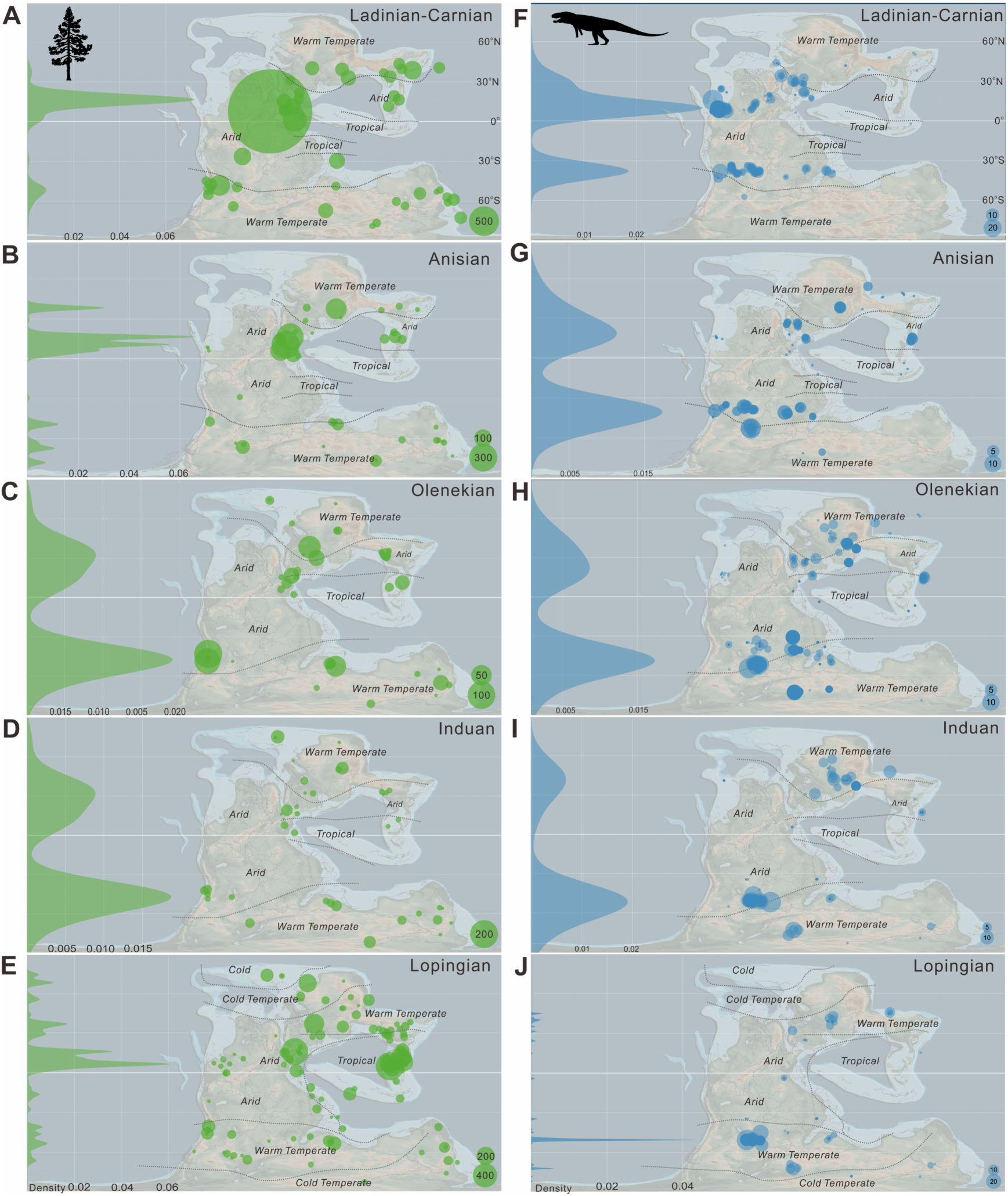
Spatio-temporal distribution of plants and vertebrates during the Permian Triassic interval. A-E. Latitudinal occurrences gradient (green curves) and heat maps (green disks) of spatio-temporal distribution of macroplants on paleogeographic maps spanning from the Lopingian (late Permian) to post-Anisian (Late Triassic). F-J. Latitudinal occurrences gradient (blue curves) and heat maps (blue disks) of spatio-temporal distribution of vertebrates. All the palaeogeographical maps are based on Scotese (2021).

Vertebrate distributions during the Lopingian are widespread across Pangaea, but are strongly concentrated in warm temperate regions (Figure 2J). Vertebrates are sparse in tropical arid belts and are largely absent from cold climatic zones. Across the PTB, vertebrate distributions become increasingly restricted. Vertebrates remain concentrated within warm temperate belts in North Hemisphere during the Induan and Olenekian (Figure 2H, J). By the Anisian, vertebrate occurrences expand modestly in both hemispheres, though distributions continued to be centred on warm temperate climates (Figure 2G). A more pronounced geographic expansion is evident by the Ladinian - Carnian, with increased occurrence in the arid zone (Figure 2F).

The latitudinal occurrences gradient (LOG) is evident in both plants and terrestrial vertebrates through the PTB. Their LOG is reorganized, and exhibits similar LOG after the PTB. Before the PTB, plant occurrences span most latitudinal intervals, with a particularly high concentration at 0-30°N in the Lopingian Epoch (Figure 2E). The maximum of the distribution occurs at the 30-60° intervals during Induan and Olenekian ages (Fig 2C, D). During the Anisian and post- Anisian, the distribution extends further, re-establishes a LOG pattern similar to that before the PTB, with fossil occurrences again concentrates at 0-30° N (Figure 2B). Vertebrate occurrences are consistently centred in the mid-latitudes during the Lopingian Epoch (Figure 2J). During the Induan, occurrences are largely confined to the 30-60° latitudinal belts in both hemispheres (Figure 2I), whereas from the Olenekian to the Carnian they are primarily distributed between 30-60°S and 0-30°N (Figure 2F-H).

The diversity of macroplants and vertebrates shows a latitudinal heterogeneity, with distinct varying responses to the EPME across different latitudinal zones. For macroplants, diversity within 0-30°N exhibits a prolonged decline of 33.3 % from late Wuchiapingian to early Olenekian (Figure 3A), followed by a relatively stable diversity value from the middle Olenekian to Ladinian (Figure S5A-C), whereas in the 0-30°S, plant diversity declines sharply by 84.38% between late Changhsingian and Olenekian, showing a more abrupt transition relatively to the low latitude in the Northern Hemisphere (Figure 3B). Plant diversity in mid latitudes (30-60°) increases progressively from the early Changhsingian through the Induan, remains relatively stable during the Olenekian and subsequently declines in the Northern Hemisphere (Figures 3A and S5D-I) and peaks at the end of Early Triassic in the Southern Hemisphere. Plant diversity in high latitudes (60-90°) declines in the Northern Hemisphere but keeps increasing in the Southern Hemisphere (Figures 3A, B and S5J-R). Vertebrate diversity exhibits pronounced fluctuations across the PTB. At the 0-30°N interval, diversity declines from the late Wuchiapingian to the Induan, reaches a minimum around the PTB, and then rises sharply to a peak in the middle Anisian before declining again (Figures 3C and S7A-C). In the 30-60°N zone, the diversity also declines from the late Wuchiapingian to the PTB, and remains low thereafter (Figures 3C and S7D-F). In the Southern Hemisphere, only 30-60° latitudinal zone is assessed. Diversity shows a sharp decline from the late Wuchiapingian to Induan. Vertebrate diversity remains stable during the Olenekian, and undergoes a subsequent rise and fall during the Anisian (Figures 3D and S7G-I). This diversity pattern closely resembles that observed at 30-60° N.

**Figure 3.**
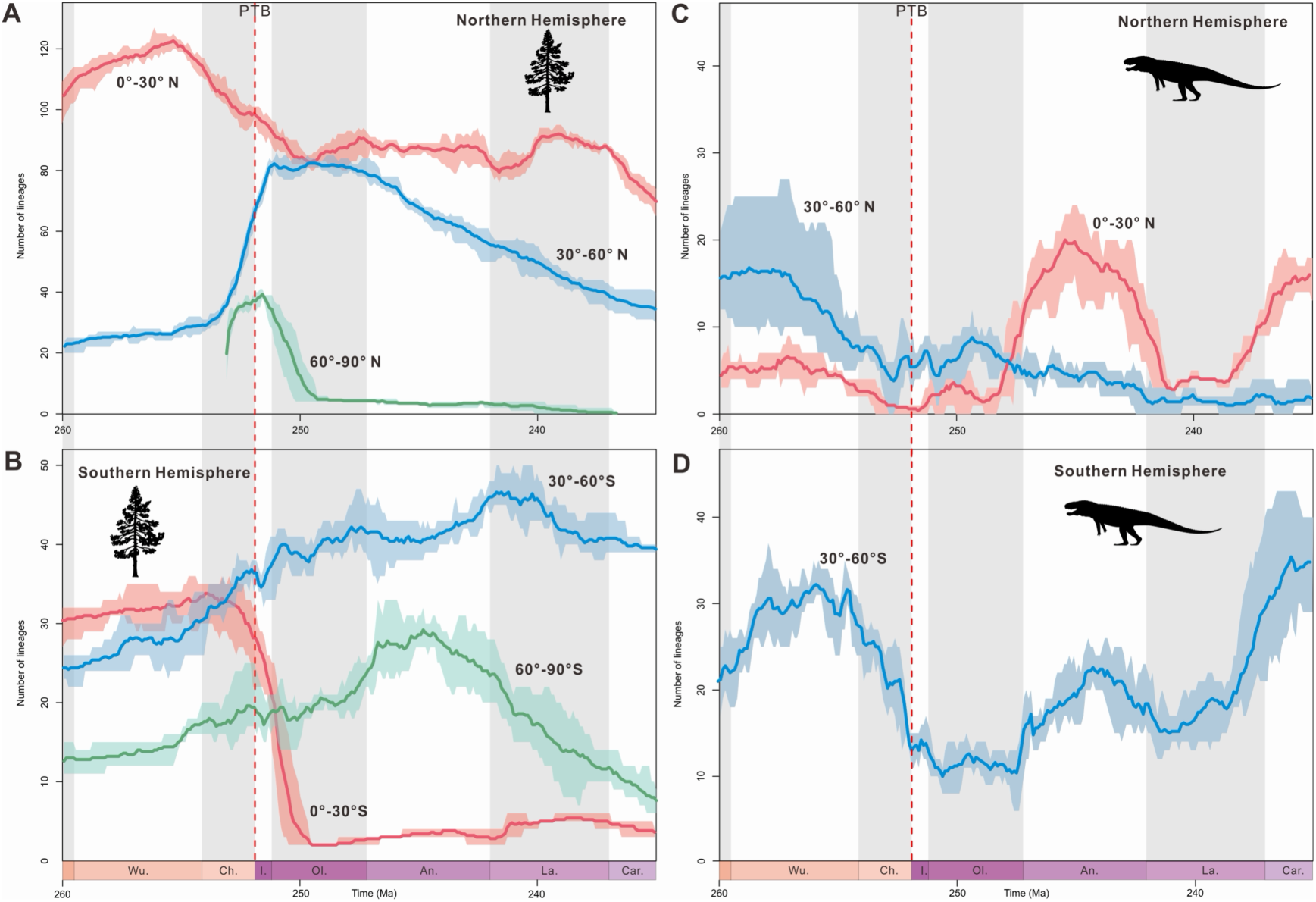
Dynamics of macroplant and vertebrate genus-level diversity at different latitudinal zones. A. B. Macroplant diversity in Northern and Southern hemispheres. The diversity in 0°-30° N shows a continuous decline from the late Wuchiapingian to middle Olenekian, and follows by a flatter trend to Carnian. 30°-60° N and 60°-90° show increase across the PTB. The diversity in 0°-30° N shows a rapid decline from Changhsingian to middle Olenekian, and 30°-60° N and 60°-90° show sustained increase. The diversity pattern of macroplant is heterogeneous in latitudinal zones. Note that both increasing and decreasing trends occur through the PTB. For origination, extinction and net diversification rates see in Figure S5. C. D. Vertebrate diversity in Northern and Southern hemispheres, only 0°-30° N, 30°-60° N, and 30°-60° S estimated. The diversity in 0°-30° N shows a decline from middle Wuchiapingian to middle Changhsingian, after which diversity remains low. The diversity in 30°-60° N shows similar pattern to 0°-30° N, with the difference that its diversity increases rapidly from late Olenekian. Note that there are no dramatic declines through or after the PTB. For origination, extinction and net diversification rates see in Figure S7. The 60°-90° N, 0°-30° S and 60°-90° S are without sufficient data to generate diversification estimates. The silhouettes were taken from https://www.phylopic.org/.

### Drivers of diversification

We test five factors that potentially drive plant diversification through the PTB using multivariate birth–death (MBD) models. Separately, we test fourteen factors that potentially drive vertebrate diversification through the PTB using the same framework (Figure 4). Most factors showing significant influence on plant and vertebrate origination and extinction (ω>0.5) are characterized by regional heterogeneity (Tables S1-18).

**Figure 4.**
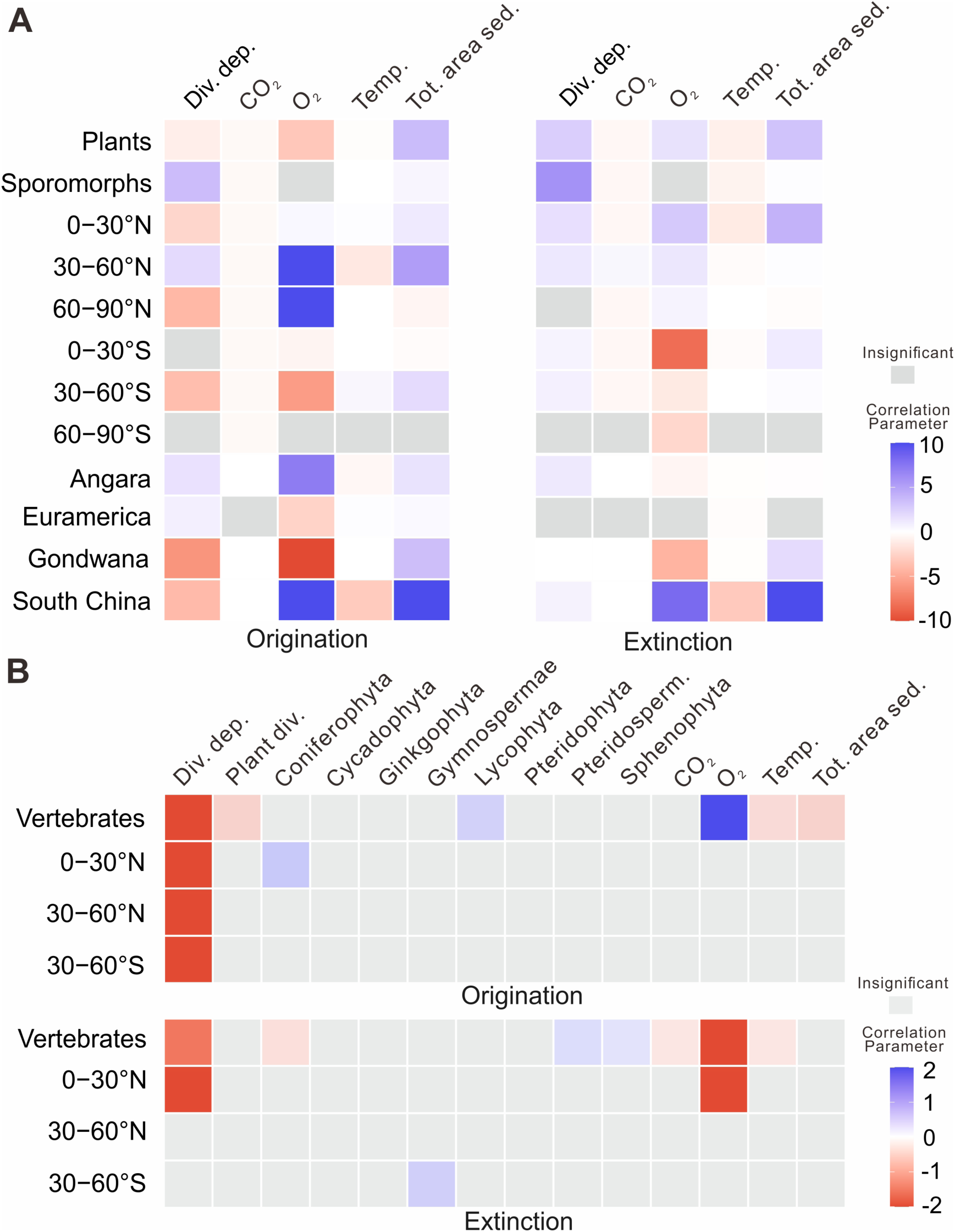
Biotic and abiotic drivers of diversification rate of macroplants (A) and terrestrial vertebrates (B) through the PTB. Correlation parameters (*Gi*) for macroplant origination and extinction. Colors indicate significant correlation (shrinkage weight > 0.5) and gray indicates the insignificant correlation (shrinkage weight < 0.5), red is negative correlation and blue is positive correlation. Div. dep.: Diversity dependence; Temp.: Temperature; Tot. area sed.: Total area covered by sediments; Plant div.: Plant diversity.

In global plant diversification, diversity dependence (*Gλ*=-0.98, *ωλ*=0.72), atmospheric O_2_ (*Gλ*=-3.23, *ωλ*=0.83) and CO_2_ (*Gλ*=-0.0004, *ωλ*=0.74), and temperature (*Gλ*=-0.16, *ωλ*=0.77) are negatively correlated with plant origination (Figure 4A and Table S1), and combine to limit plant diversification. Total area covered by sediment is the only factor positively correlated with origination (*Gλ*=3.63, *ωλ*=0.97), suggesting that the habitat availability or high preservation rate may have facilitated the observable signal of plant diversification. Temperature (*ωµ*=0.99) and total area covered by sediment (*ωµ*=0.95) are again associated with high shrinkage weights. Temperature has a substantial negative effect to extinction (*Gµ*=−0.86), indicating that cooler condition significantly increases plant extinction risk. Total area covered by sediment again has a positive effect to origination (*Gµ*=3.24) (Figure 4A and Table S1). Interestingly, total area covered by sediment exerts a strong positive influence on both origination and extinction rates of global plant diversity.

For sporomorphs, diversity dependence shows a strong positive correlation with both origination (*Gλ*=3.52, *ωλ*=0.87) and extinction (*Gμ*=5.93, *ωµ*=0.95) (Figure 4A and Table S2), indicating a dynamic turnover pattern in sporomorph diversity. Temperature exerts a negative correlation with extinction (*Gμ*=-0.66, *ωµ*=0.98) (Table S2).

For the global vertebrate diversification, diversity dependence, plant diversity, temperature and total area covered by sediment are strong negative correlations with origination (*ω*>0.5) (Figure 4B and Table S15). Lycophyta diversity and O_2_ level show positive correlations with origination (*ω*>0.5) (Figure 4B and Table S15). Diversity dependence, Coniferophyta diversity, CO_2_ and O_2_ level and temperature have negative effect on extinction, Pteridospermatophyta and Sphenophyta show negative correlations with extinction (Figure 4B and Table S15). For vertebrates at 0-30°N, diversity dependence is negatively associated with origination (*Gλ*=-2.55, *ωλ*=0.75), and Coniferophyta diversity shows positive correlations with origination; Diversity dependence and O_2_ level have negative effects on extinction (Table S16). Only diversity dependence shows strong negative correlations with origination in 30-60°N (*Gλ*=-4.29, *ωλ*=0.89) (Figure 4B). In 30-60°S, diversity dependence and Gymnospermae diversity have a negative effect on origination (*Gλ*=-2.04, *ωλ*=0.73) and positive on extinction (*Gµ*=0.08, *ωµ*=0.53), respectively.

## DISCUSSION

### Diversification trends of terrestrial organisms through the PTB

Macroevolutionary patterns among Phanerozoic biotic crises range from collapse-dominated mass extinctions to expansion-driven radiations (Hoyal Cuthill et al., 2020). Classic ones, such as EPME (252 in Figure 5B) and End-Devonian crisis (382 in Figure 5B), are characterized by rapid biodiversity loss (high extinction rate) coupled with delayed ecological recovery (low origination rate). In contrast, Permo-Triassic terrestrial plants and vertebrates are stand on markedly different position within this framework, being closer to an extinction-radiation domain than to a genuine extinction regime (Figure 5B).

**Figure 5.**
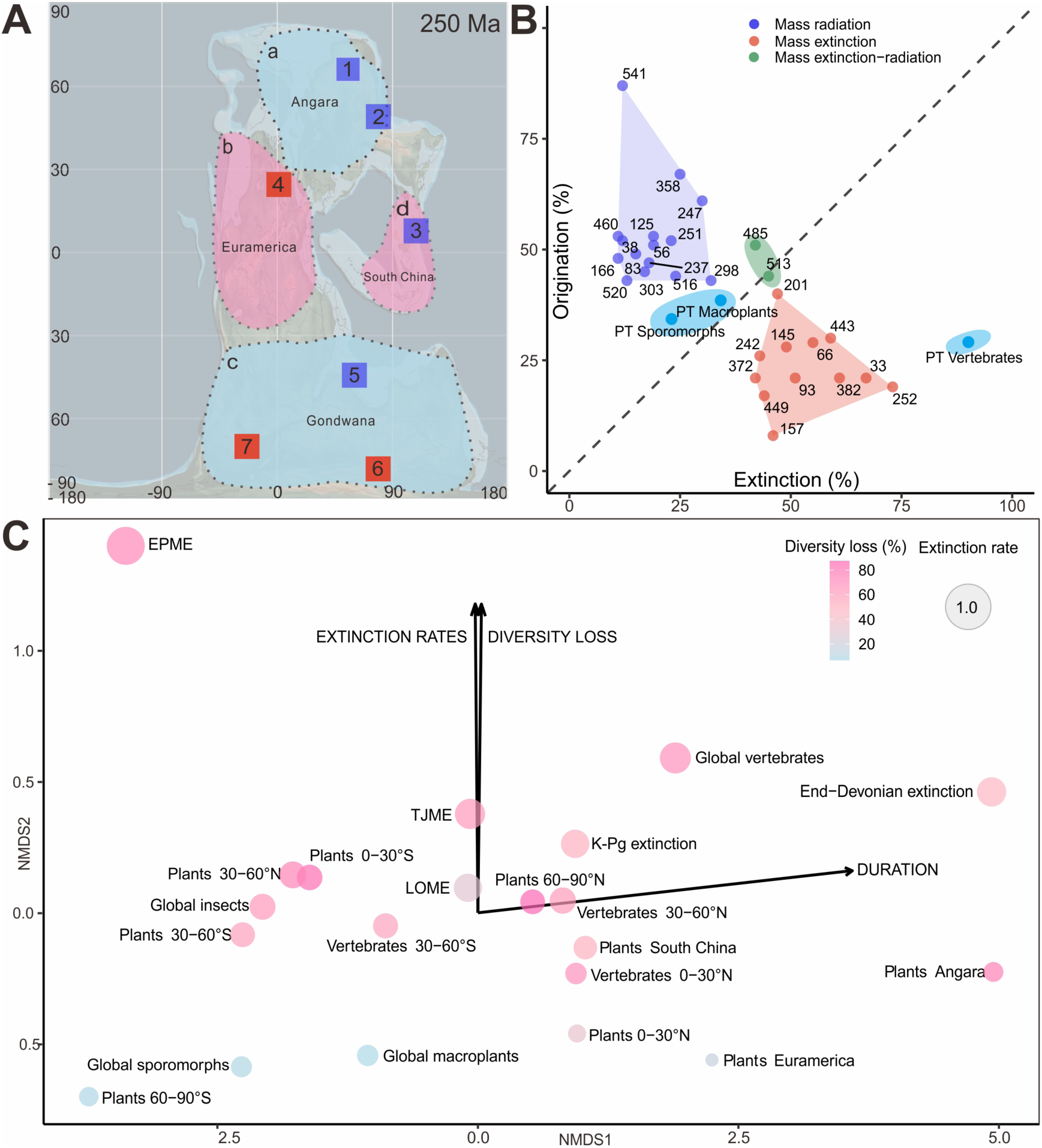
Macroevolution evidenced by plant biogeography, major biotic events, and Non-metric multidimensional scaling (NMDS) analysis during the PTB. **A** Extinction in phytogeographic regional scale on late Permian paleogeographic map. A (Angara) and C (Gondwana) regions (light blue) do not exhibit a significant decline in diversity, and B (Euramerica) and D (South China) regions (light red) show a marked decrease in diversity. Individual fossil specimen-based study are indicated by locality numbers 1-7 1, Kuznetsk Basin, Siberia, Russia, and the diversity of terrestrial organisms not showing a significant decline (Davydov et al., 2021); 2, Junggar Basin, China, with a diverse terrestrial ecosystem spanning the PTB (Peng et al., 2025); 3, Meishan Section, South China, its palynological evidence showing a gymnosperm-dominated vegetation that thrived through the PTB (Schneebeli-Hermann and Galasso, 2025); 4, East Greenland, its palynological evidence revealing strongly time-delayed end-Permian terrestrial plant extinctions, with a transient increase in plant diversity (Looy et al., 2001); 5, southern Tibet, China, the EPME with only short-term disturbance in terrestrial vegetation based on palynology (Liu et al., 2020); 6, Sydney Basin, Australia, significant impact of EPME on plants (Mays et al., 2021); 7, Karoo Basin, South Africa, vertebrates with a high regional extinction rate within∼ 1 Myr (Viglietti et al., 2021). Palaeogeographical map is based on Scotese (2021). **B** Three types of relationship between extinction and origination across major biotic events: mass radiation, mass extinction and mass extinction-mass radiation. Numbers indicate the time of the events (Ma). The PT data is based on our calculations, others from Hoyal Cuthill et al. (2020). **C** Non-metric multidimensional scaling (NMDS) ordination of extinction dynamics across major Phanerozoic biotic crises. The distribution of plants and vertebrates across palaeolatitudinal regions and phytogeographical zones reveals strong spatial and taxonomic heterogeneities, highlighting fundamental differences in tempo and magnitude of terrestrial dynamics during the PTB. Bubble size indicates extinction rates and colour denotes percentage of diversity loss, while vectors summarize the dominant gradients of extinction rates, diversity loss and duration. Data of marine extinction event are based on publications. LOME, late Ordovician mass extinction; TJME, Triassic-Jurassic mass extinction; K-Pg extinction, Cretaceous-Paleogene extinction event.

Macroplants and sporomorphs across the PTB exhibit moderate extinction and substantial origination, indicating that land plants of this period underwent turnover and reorganization instead of global biotic collapse. Global flora shows lower extinction rates and lower extinction rates than other terrestrial clades across the PTB, and no tremendous, large-area, or mass extinction in light of macroevolution concept (Raup et al., 1982; Hull et al., 2015). Vertebrates show a pronounced global decline in diversity spanning ∼6 Myr, reflecting a gradual, long-term decay rather than an instantaneous event. The decoupling between extinction intensity and origination in terrestrial systems suggests that land ecosystems responded to Permo-Triassic environmental perturbations through restructuring of community composition and ecological strategies, rather than simple diversity bottlenecks.

### Diversity across latitude and regions

Extinction dynamics across the PTB exhibit pronounced heterogeneity among groups and latitudinal and regional gradients (Figure 5C). The extinction rates of plants, including macroplants and sporomorphs, show only modest increases during the EPME and remain comparatively low to those of terrestrial vertebrates (Figure 5C), which, in contrast, show display strongly high extinction rates, vertebrates reaching a maximum extinction rate of 1.1022, the highest value among all terrestrial clades (Figure 5C and Table S19). Such contrast is indicated by a clear separation between plant and vertebrate along the extinction-rate gradient in our NMDS ordination (Figure 5C).

Latitudinal patterns further reveal spatial structuring of extinction and diversity loss. Macroplants exhibit high extinction rates primarily in the 0-30°S and 60-90°N intervals, whereas in other latitudinal belts, close to background levels. Plant diversity declines sharply at low latitudes, the loss reaching 84.38% in the 0-30°S interval and 33.3% at 0-30°N, while in mid-latitude regions (30-60°N), stable or even increased diversity across the PTB. Such latitudinal asymmetry is consistent with the hypothesis that collapse of tropical vegetation exerted a disproportionate influence on Earth system dynamics during the EPME. Severe diversity losses in tropical and subtropical regions probably amplify environmental instability (Xu et al., 2024; Chen et al., 2025), contributing to prolonged post-extinction perturbations. In contrast, the stability of diversification among mid- to high-latitude plants suggests more stable environment or plants in this latitude with higher resilience.

Vertebrate diversity also shows pronounced latitudinal heterogeneity. Diversity in the 0-30°N interval declines sharply during the end-Permian, approaching near-collapse. Mid-latitude vertebrate assemblages (30-60°N and 30-60°S) exhibit higher diversity than equatorial regions, forming a bimodal latitudinal distribution that contrasts with the modern latitudinal diversity gradient. These observations might indicate a potential equatorial tetrapod gap or escape from the equatorial (Bernardi et al., 2018; Allen et al., 2020; Liu et al., 2022). In the NMDS ordination, vertebrate datasets plot toward higher extinction-rate vectors, reinforcing the view that terrestrial vertebrates experienced more intense and less buffered extinction dynamics than plants (Figure 5C). Nonetheless, the incompleteness of the vertebrate fossil record in some regions highlights the need for further data to fully resolve these patterns.

Regional heterogeneity in plant diversity is further supported by biome-level analyses. Nowak et al., (2020) assigned fossil plants to different biomes and indicated profound but heterogeneous changes in terrestrial vegetation, with a marked reduction in biome diversity and a shift toward increased seasonality following the PTB. Our study demonstrates that the disappearance of humid-tropical and cool-temperate floras after the PTB (Nowak et al., 2020) coincides with the significant diversity loss observed from the South China (tropical area) and Angara (cold and warm temperate) floras, and that the reduction of tropical and cool-temperate biomes probably contributes to the Early-Middle Triassic “coal gap” (Nowak et al., 2020).

Summarizing the above, it can be concluded that terrestrial diversity across the PTB is spatially heterogeneous and fundamentally decoupled with overall trends, underlying the regional and ecological roles in shaping extinction dynamics.

It is noted that preservation biases inherent to terrestrial organisms represent a fundamental constraint (Alory et al., 2008; Servais et al., 2019; 2023; Capel et al., 2022). Fossil specimens-based study is prone to be influenced by such biases, which are hardly eliminated even by means of different analytical approaches (Silvestro et al., 2014; 2019). In this dataset, fossil occurrences of vertebrates in 0-30°S and 60-90°S remain particularly sparse or absent, preventing reliable model fitting in these regions.

Meanwhile, the apparent dual effect of sedimentary area may, at least in part, reflect preservation and sampling biases rather than purely biological signals. More and continued accumulations of fossil discoveries are essential for improving our understanding of biodiversity dynamics in insufficient sampling areas, and of course will minimize the limitation of or similar to our study.

### Heterogeneous response under global drivers

Our multivariate birth-death analyses demonstrate that diversification across the PTB shows strongly heterogeneous with regional and taxonomic responses to environment stress, lacking of uniform global drivers. Multiple abiotic and biotic factors influenced origination and extinction rates, but their direction and intensity vary markedly among plants, sporomorphs and vertebrates, and across latitudinal zones (Figure 4).

Globally, plant diversification appears to have been constrained by interacting ecological and physiological limits. Negative associations between plant origination and diversity dependence, temperature, atmospheric CO_2_ and O_2_ suggest that ecological saturation and climatic stress jointly suppressed the emergence of new lineages during the PTB. In contrast, sedimentary area shows a consistent positive association with both origination and extinction, highlighting its dual role in promoting diversification through habitat expansion while simultaneously increasing extinction risk through environmental instability.

Vertebrate diversification exhibits a different dependency structure. Negative effects of diversity dependence and temperature on origination suggest that vertebrate diversification was constrained when ecosystems became crowded or when thermal conditions were unfavorable. The positive associations with Lycophyta diversity and O_2_ indicate that vertebrate diversification was promoted under conditions of higher primary productivity and improved physiological support. These patterns suggest that vertebrate diversification during the PTB was influenced more by the structure and productivity of terrestrial ecosystems, and by oxygen availability, than by climate alone.

The influence of these drivers is spatially heterogeneous. Temperature and CO_2_ exert particularly strong effects in low-latitude regions, whereas CO_2_ shows contrasting impacts at higher southern latitudes, promoting origination in the 30-60°S zone but inhibiting it in the 60-90°S interval. Such spatially variable responses underscore that diversification dynamics were regionally modulated, with identical global drivers producing divergent evolutionary outcomes depending on local environmental thresholds and ecological structure. These results demonstrate that diversification across the PTB is not uniformly controlled by global environmental drivers. This spatially heterogeneous response provides a mechanistic framework for understanding the pronounced regional differences in terrestrial extinction and recovery patterns observed across the Permian-Triassic transition.

While we explored the influence of multiple biotic and abiotic drivers on biodiversity dynamics and examined how different latitudinal bands and regions respond to global-scale drivers, this approach entails important interpretative and data-related constraints. The availability and temporal resolution of environmental reconstructions vary substantially across regions and latitudes (Jones and Eichenseer, 2021), limiting our ability to further subdivide environmental variables and to investigate finer-scale evolutionary patterns. Moreover, environmental drivers were implemented at a global scale, whereas their biological effects are inherently latitude-dependent. As a result, global mean variables, such as temperature or continental fragmentation, may reflect contrasting selective regimes across latitudes, potentially obscuring regionally specific evolutionary responses. At present, the limited availability of high-resolution, latitude and region-specific environmental data prevent a fully spatially explicit treatment of explanatory variables.

Beyond environmental drivers, we also used the DES framework to investigate latitudinal dispersal of plant and vertebrate, and its potential influence on regional extinction dynamics. Although inter-latitudinal dispersal was persistent across the Permian-Triassic interval, it did not intensify during the crisis and shows no clear association with elevated extinction rates. While dispersal likely modulated regional diversity patterns, it does not appear to have been the primary driver of terrestrial biodiversity restructuring, which was instead dominated by climatic and other compounded environmental stresses (Figures S6, 8, 10).

### Mosaic diversification pattern

We demonstrate that terrestrial organisms (plants and vertebrates) during the PTB do not undergo a global and synchronous mass extinction, instead only the diversity crisis or prolonged interval of spatially heterogeneous ecosystem turnover, with extinction and origination being tightly coupled. We summarize terrestrial diversification through the PTB as the mosaic diversification pattern, referring to the spatio-temporal and taxonomic heterogeneities of diversity dynamics, in which origination and extinction show no uniform recovery or collapse but are driven by a complex interplay of distinct regional and latitudinal factors. Global diversification is not shaped by monolithic drivers, but by a patchwork of localized dynamics, akin to pieces of a mosaic, that collectively form the broader evolutionary dynamics.

Accumulating fossil evidence independently supports a diachronic and spatially heterogeneous model of terrestrial diversification across the PTB. As in South China at low latitudes show a clear extinction event and a significant decrease in fossil occurrences and diversity (Figure 5A) (Feng et al., 2020; Xu et al., 2022), whereas in the coeval Euramerica and Gondwana regions no significant decline is seen (Figure 5A). In the Sydney Basin, the disappearance of *Glossopteris* flora occurred prior to 252.3 Ma, causing by the onset of Siberian Traps Large Igneous Province (Fielding et al., 2019).

Palynological evidence shows only temporary disturbances and range reductions of terrestrial plant communities of southern Tibet in EPME (Liu et al., 2020). In Karoo Basin, South Africa, vertebrate turnover predates the marine mass extinction and does not represent the terrestrial end-Permian mass extinction (Gastaldo et al., 2015; 2020). The plant assemblage from Permian equatorial lowlands reveals that three major plant clades (Bennettitales, Corystospermales, and Podocarpaceae) cross the PTB, were less severely affected by global biotic crises than traditionally assumed (Blomenkemper et al., 2018). Gymnospermous forests and ferns persisted across PTB, and did not suffer from extinction in Junggar Basin (Peng et al., 2025) and sporomorph records from the Global Stratotype Section and Point of the PTB in the Meishan Section indicate that no significant floral changes and gymnosperms consistently dominate throughout the PTB (Schneebeli-Hermann and Galasso, 2025) (Figure 5A).

### Data and code availability

All data are available in the manuscript or in the supplementary information. All code used in this analysis, along with all necessary files, are available on Figshare (…..).

## ACKNOWLEDGMENTS

We thank Ms. Yang Ning (Nanjing Institute of Geology and Palaeontology, Chinese Academy of Sciences) for data collection. This research was supported by the National Key R&D Program of China (2022YFF0800200), National Natural Science Foundation of China (No. 42202007), Shandong Provincial Natural Science Foundation (ZR2022QC257), and State Key Laboratory of Palaeobiology and Stratigraphy (NIGP, CAS) (No. 223127).

## AUTHOR CONTRIBUTIONS

Conceptualization: H.X. Methodology: B.L. H.X. Investigation: H.X. B.L. K.W.

Y.W. Visualization: B.L. K.W. Y.W. Supervision: H.X. Writing-original draft: B.L.

H.X. Writing-review & editing: B.L. H.X.

## MATERIALS AND METHODS

### Fossil occurrence data

The compilation of the dataset is based on previous dataset and we added some new fossil occurrences data. Global macroplant and sporomorph datasets are based on Nowak et al. (2019), terrestrial vertebrate is based Button et al. (2017). We removed the synonyms, sp. and cf. and divided the dataset by latitudes, classification and phytogeographical regions to further analysis. The final dataset contains 12,936 macroplant occurrences, 52,486 sporomorph occurrences and 1,609 terrestrial vertebrate occurrences.

### Occurrences data analysis

GPS coordinates for the fossil localities were obtained from the original publications or extracted from Google maps. The PaleoGPS data were calculated using the PALEOMAP model of rgplates package (Müller et al., 2018; Kocsis et al., 2024) within software R (v4.4.1). The frequency of occurrence for each locality was calculated and plotted heatmap using ggplot2 package (Wickham and Sievert, 2009), where the size of the circles represents the number of fossil occurrences. The heatmap was plotted on the paleogeographic maps (Scotese 2021; http://www.scotese.com/) in different time bins. We combined the paleoclimatic zones based on the Boucot et al., 2013 on paleogeographic maps. The heatmaps show the fossil occurrence density of different regions and the distribution patterns fossil with changing climatic zones.

### Estimation of diversification

We analyzed the global plant and vertebrate occurrences datasets, and different latitude zones datasets using the PyRate program (Silvestro et al., 2014; 2014b; 2019) under the Reversible Jump Markov Chain Monte Carlo (RJMCMC) algorithm, which provides more accurate results, to investigate the speciation (*Ts*) and extinction (*Te*) times of each lineage, origination and extinction rates through time, and net diversification rate. The best-fitting preservation model was tested using the *-PPmodeltes*t option, and Non-homogeneous Poisson process of preservation (NHPP) Homogeneous Poisson process (HPP) and Time-variable Poisson process (TPP) were assessed as being best supported by the data. TPP model was identified the best-fitting preservation model, and all datasets were calculated using TPP model.

Each Markov Chain Monte Carlo (MCMC) run executed for 30,000,000 generations, sampling every 30,000 generations, and were replicated across five independent chains (*-j1* to *-j5*) to ensure convergence of every dataset. Posterior samples from the independent MCMC runs were combined using *- combLogRJ*, discarding the first 10% as burn-in (-*b 0.1*), and visualized the rate variation through time in genus- and species-level of global plant and animal. Then, the output files were used to produce lineage through time (LTT) plot based on range-through diversity (*-ltt* option). Here, we estimate terrestrial diversity in 0.1 Myr time bins, derived from the calculated *Ts* and *Te* of each lineage, allowing for a high-resolution depiction of diversity patterns and to understand details of diversity dynamics. All the log files were checked in Tracer v1.7.2 (Lehtonen et al., 2017) to estimate the effective sample size and convergence of parameters sufficient.

### Multivariate birth-death model

We used the multivariate birth-death model (MBD) to asses to what extent biotic and abiotic factors can explain temporal variation in origination and extinction rates, and calculated in PyRateMBD (61). In the MBD model, origination and extinction rates can change through time through with time-continuous variables. The strength and sign (positive and negative) of the correlations are jointly estimated the baseline origination (*λ0*) and extinction (*µ0*) rates and all correlation parameters (*Gλ* and *Gµ*) using a horseshoe prior to control for overparameterization and for the potential effects of multiple testing using the MCMC. In this model, shrinkage weights (*ω*) provide a robust measure to distinguish between noise and signal, shrinkage weights > 0.5 indicate significant support for the corresponding correlation parameter, and shrinkage weights < 0.5 indicate insignificant support.

For the plant diversification, five factors are identified to measure their impact to plant diversity at the genus-level. Diversity dependence is estimated to measure the effect on its own origination and extinction rates. Atmospheric CO_2_ and O_2_ concentrations, temperature and total area covered by sediment are regarded as significant abiotic factors for plant diversity (Salles et al., 2023). In the terrestrial vertebrate diversification, 15 factors are identified to estimate the effect to animal diversity in genus-level. Diversity dependence and other 8 biotic factors including total plant diversity, and each major plant group diversity, Coniferophyta, Cycadophyta, Ginkgophyta, Gymnospermae *incertae sedis*, Lycophyta, Pteridophyta, Pteridospermatophyta and Sphenophyta, as well as four abiotic factors are estimated, containing atmospheric CO_2_ and O_2_ concentrations, temperature and total area covered by sediment. Data of total plant and each major plant group diversities based on the calculation in PyRate, CO_2_ from Foster et al. (2017), O_2_ from Lehtonen et al., 2017, temperature from Prokoph et al. (2008) and total area covered by sediment based on the data from Salles et al., 2023.

We ran the MBD model using 100,000,000 MCMC iterations and sampling every 20,000 generations to approximate the posterior distribution of all parameters based on the log files obtained from previous PyRate TPP model analysis. The default model used in MBD analysis was exponential correlations (-*m0*). The time interval analyzed ranged from 264 to 227 Ma across the Permian-Triassic transition. The results of MBD were summarized by calculating the posterior mean and 95% HPD of all correlation parameters (*Gλ* and *Gµ*) and the mean of the respective shrinkage weights (*ω*), and the mean and 95% HPD of the baseline origination and extinction rates (*λ0* and *µ0*). The plant data was divided to global plant and spores, 6 latitude zones, 0−30°N, 30−60°N, 60−90°N, 0−30°S, 30−60°S, 60−90°S, and we performed MBD analysis on these 10 datasets. The vertebrate data was divided to global data and 0−30°N, 30−60°N, 30−60°S latitude zones, due to the lack of data in other latitude zones, calculations were not performed. The division of latitudinal interval is based on calculate paleolocations. All the log files were inspected in Tracer v1.7.2 (Drummond and Rambaut, 2007) to check the effective sample size and convergence of parameters sufficient.

In this study, we calculated origination, extinction, net diversification rates and diversity within the time interval of 266.9-205.7 Ma (Wordian to Norian). Because fossil records are relatively sparse at the temporal margins, the curves at both ends of the interval may be affected by undersampling (edge effects). To provide a clearer assessment of terrestrial diversity changes across the PTB, we restricted the plotted results to 260-245 Ma, thereby focusing on intervals with comparatively richer fossil records and more robust estimates. All the calculations in PyRate were done from 10/2024 to 5/2025, we used numerical age in the International Chronostratigraphic Chart v2023/06 to measure the origination, extinction and diversification.

### Dispersal Extinction Sampling models

Diversity dynamics in plants and animals are influenced by dispersal among different regions (Carrillo et al., 2020; Nowak et al., 2022; Liu et al., 2024). To quantify spatiotemporal variation in dispersal and extinction dynamics of terrestrial plants and vertebrates across the Permian-Triassic transition, we applied the Dispersal Extinction Sampling (DES) model implemented in PyRate (Carrillo et al., 2020). Fossil occurrence data were compiled separately for plants and vertebrates and subdivided into multiple latitudinal bins to explicitly test geographic heterogeneity in macroevolutionary dynamics. All the plant and vertebrate occurrence data were assigned to 0−30°N, 30−60°N, 60−90°N, 0−30°S, 30−60°S, 60−90°S, according to the paleolatitude. Then, DES model was used to calculate the dispersal and extinction rates between adjacent latitudinal zones.

Here, we used time variable model (*-TdD, -TdE*) with rate shifts (Skyline model) in DES allowing rate to shift across predefined temporal boundaries spanning the Late Permian to Late Triassic (266.9-205.7 Ma). Fossil age uncertainty was accommodated through replicated datasets generated using a fixed temporal bin size of 0.5 Myr. Markov Chain Monte Carlo (MCMC) were run 500,000 iterations, sampling every 1,000 steps, with an initial burn-in removed prior to inference. Multiple independent replicates were performed for each latitudinal bin to assess convergence and robustness of rate estimates. Posterior samples from replicate runs were combined to obtain marginal rate-through-time trajectories and associated credibility intervals. Identical analytical pipelines were applied to plants and vertebrates, enabling direct comparison of dispersal and extinction dynamics across taxonomic groups and latitudinal zones (Figure S1).

