## Supplemental Information for "Mosaic terrestrial diversity dynamics through the Permo-Triassic interval"

**About this research**

This study investigates the spatio-temporal dynamics of terrestrial organisms across the Permo-Triassic boundary (PTB) by integrating large-scaled fossil occurrence data with quantitative diversification and biogeographic modelling frameworks (Figure S1). Our overarching aim is to precisely depict how extinction, origination, dispersal and sampling processes jointly shaped terrestrial biodiversity patterns during the end-Permian mass extinction (EPME), one of the most severe biotic crises in geologic history.

We compiled and curated a comprehensive global dataset of fossil occurrences spanning from the late Permian to Middle Triassic, encompassing three major terrestrial biotic groups: macroplants, sporomorphs, and vertebrates. After taxonomic standardization, temporal calibration and data cleaning, the final dataset includes occurrences of 12,936 macroplants, 52,486 sporomorphs, and 1,609 vertebrates. These organisms consist of different ecologic strategies, fossil preservation potential and spatial signal, and allows comprehensive and direct comparison of diversity dynamics (Figure S1A).

To quantify diversification dynamics through time, we applied a Bayesian birth-death framework implemented in *PyRate*, using reversible-jump Markov chain Monte Carlo (RJMCMC) algorithm. This approach jointly estimates origination rates, extinction rates, diversity trajectories and preservation rates while explicitly accounting for heterogeneous sampling. Analyses were conducted at multiple analytical resolutions, including global genus-level and species-level datasets, as well as subsets partitioned by latitude intervals, biogeographical regions and ecological groupings. This hierarchical design enables assessment of both global trends and regionally structured responses across the Permo-Triassic transition.

Spatio-temporal distribution patterns were reconstructed using paleogeographic projections based on the PALEOMAP model implemented through the *rgplates* framework. Fossil occurrences were rotated into paleocoordinate space to evaluate changes in latitudinal distribution, biogeographical range shifts and regional occupancy through time. These reconstructions provide a spatial context for interpreting diversification dynamics and enable direct comparison between biological turnover and paleogeographic reorganization.

To further explore potential drivers of diversity change, we implemented a multivariate birth-death (MBD) modelling framework that incorporates both biotic and abiotic predictors. This analysis assesses the extent to which diversification rates covary with environmental variables and ecological factors, offering insights into mechanisms underlying observed diversity patterns. In parallel, dispersal extinction sampling (DES) models were applied to evaluate latitudinal differences in dispersal dynamics and extinction intensity, explicitly accounting for uneven sampling across regions.

By integrating diversification modelling, paleogeographic reconstruction and dispersal analyses within a unified workflow, this study provides a quantitative framework for linking temporal diversity dynamics with spatial reorganization of terrestrial ecosystems across the PTB. The approach highlights how differential extinction, recovery trajectories and geographic restructuring collectively shaped the evolution of terrestrial life during this critical interval.

**Origination and Extinction in genus level**

In this study, we estimated the extinction and origination rates using PyRate framework. As a complementary measure to these rate estimates, we also calculated the percentage of origination and extinction from the Changhsingian to Induan ages based on the genus duration (Figure S4). The calculation of origination and extinction are based on

$$Origination\ \% = \frac{N_{originated}}{N_{pre}} \times 100\%$$

$Extinction\ \% = \frac{N_{extinct}}{N_{pre}} \times 100\%$ (Figure S4).

The results indicate that macroplants experiences an extinction rate of 33.3% and an origination rate of 39.4%, sporomorphs show 22.2% extinction and 35.2% origination, whereas vertebrates suffer extremely high extinction (90.1%) coupled with much lower origination (29.1%) from the Changhsingian to the Induan ages (Figure S4). From the perspective of origination-extinction balance across the PTB, plant (macroplant and sporomorph) groups exhibit broadly comparable rates of extinction and origination, indicating that this interval was characterized not by a global-scaled collapse of terrestrial floras but by pronounced turnover. In contrast, vertebrates experience a genuine mass extinction, marked by overwhelming losses that are not offset by contemporaneous origination.

**Diversity at four phytogeographic zones**

Four phytogeographic zones, Angara, Euramerica, Gondwana and South China, are considered to investigate the regional diversity trends. In Angara, plant diversity shows a steep rise from early Changhsingian to Induan, followed by an abrupt and sustained decline throughout the Early Triassic (Figure S9A-C). Euramerica presents a similar pre-extinction buildup in diversity, but with a less abrupt decline (16.70 % diversity loss) across the PTB (Figure S9D-F). Gondwana displays relatively muted diversity changes (Figure S9G-I). South China experiences a sharp pre-boundary diversity peak followed by a near-complete collapse of diversity, showing two pulses of decline in early Changhsingian (34.4% diversity loss) and post PTB (47.5 % diversity loss) (Figure S9J-L). Post-extinction recovery is minimal, and lineage richness remained low throughout the Early and Middle Triassic in South China paleoblock.

**Improving the resolution of terrestrial fossil and stratum records**

With accumulation of fossil records and application of new methods, temporal resolution of biodiversity has markedly improved, from the stage-level to 11 Ma time bin (Alroy et al., 2008), even to 26 ka or 21 ka (Fan et al., 2020; Deng et al., 2021). These advances have enhanced our understanding of detailed biodiversity dynamics in deep time. Notably, diversity patterns vary substantially depending on the temporal resolution (see Figure 3 in Fan et al., 2020). High-resolution diversity curves benefit from the continuity of marine strata and abundance of marine fossil records. In contrast, terrestrial strata lack continuity, making it challenging to achieve comparable resolution. Therefore, methods are crucial for improving the temporal resolution of terrestrial biodiversity estimates. The Bayesian framework (PyRate) shows accuracy and robustness in calculating datasets with various degrees of data incompleteness (Silvestro et al., 2014; 2014b; 2019), reducing the impact of incomplete fossil records and estimating diversity in different time bin. With the application of Bayesian frameworks, new insights have been gained into diversity of dinosaurs (Condamine et al., 2021), insects (Jouault et al., 2022; 2024), mammalians (Buffan et al., 2025), and angiosperms (Condamine et al., 2020). Studies of terrestrial diversity have assessed diversity primarily at the stage level (Benton and Newell, 2014; Nowak et al., 2019), limiting the resolution of diversity dynamics. In this study, we examine terrestrial diversity across the PTB and provide a more detailed characterization of diversity, capturing finer-scale patterns across clades and regions, thereby complementing global diversity reconstructions that may overlook such heterogeneity.

Preservation bias and incompleteness of fossil records are essential issues that need to be considered in diversity studies (Barrett et al., 2009; Close et al., 2020). In the terrestrial fossil records, preservation bias is evident. In this study, the scarcity or absence of plant fossil records in the 60-90°N interval, as well as the vertebrate fossil records in the 60-90°N, 0-30°S, and 60-90°S intervals, clearly reflect strong preservation bias. Such large-scale gaps are difficult to overcome through methodological advances alone. Assessing the impact of preservation rates is also crucial for reconstructing diversity in regions or zones with abundant fossil records. PyRate can calculate the preservation rates per lineage in stage level. Our results reveal that although preservation rates do affect observed diversification, especially in certain areas and groups, it does not completely obscure the real diversity signal. Macroplant and vertebrate diversity trends are decoupled from preservation rates, especially across the PTB, indicating that the diversification rate reflects the genuine dynamics in geologic history. South China and Angara show correlation between preservation rates and diversity, implying localized preservation effects. This geographical heterogeneity highlights the importance of regional assessments when explaining global diversity patterns. Overall, despite the influence of preservation rates, the trajectories of macroplant and vertebrate diversity of our analysis clades, regions and zones during the EPME reflect real evolutionary dynamics (Figures S12-15).

**Preservation rate**

Here, we simulate the preservation rates in macroplant, sporomorph, vertebrate and different latitudinal zones and areas (Figures S12-15) to explore the influence of preservation bias on diversity.

In a global scale, the preservation rates of macroplant remain high and stable throughout the late Permian, with values from 6-8, after the PTB, rates reach the peak in the Induan, following an abrupt decline in the Olenekian (Figure S12). Macroplant diversification declines in the Wuchiapingian-Changhsingian boundary and Induan, while preservation rates increase, indicating that diversification is not showing a synchronous trend with preservation rates throughout the PTB (Figure S12A). The preservation rates of sporomorph are higher than that of macroplant indicating a better preservation potential than macroplant and show similar trends to macroplant (Figure S12B). Vertebrate preservation rates show stable, below 2, during the late Permian and increase in the Induan, and post-PTB, preservation rates drop to below 2. Vertebrate diversification declines in the Wuchiapingian, while preservation rates increase; similarly, diversification initially increases and then declines, whereas preservation rates decrease (Figure S12C).

In 0-30° N zone, the preservation rates of macroplant remain high during the late Permian, more than 5, and decline in the Early Triassic (Figure S13A). Meanwhile, their diversification continues to decline from the late Wuchiapingian to middle Olenekian, inconsistent with the trends of preservation rates. In 30-60° N, preservation rates remain stable across PTB, with a temporary increase in the Induan (Figure S13B). Diversification increases form the Changhsingian to Induan, differing from the preservation rates trend. In 60-90° N, diversification declines after the PTB, showing the same trends as preservation rates (Figure S13C), indicating that the diversity in this zone is affected by preservation bias. In 0-30° S, the preservation rates of macroplant keep high rates during the late Permian, then decline in the Induan and remain at low rates thereafter (Figure S13D). Diversification is not affected by the preservation rates prior to PTB, but diversity declines as the preservation rates decline post-PTB. In 30-60° S and 60-90° S, the preservation rates also show stability in the late Permian and an increase in the Induan, and diversification does not show the similar trends to preservation rates (Figure S13E, F).

The preservation rates of vertebrate remain stable during the late Permian and decline in the Early Triassic, and then rise in the Middle Triassic in 0-30° S (Figure S14A), its diversification shows a different variation from the preservation rates. In 30-60° N and 30-60° S, the preservation rates show similar trends, diversification also fail to show any correlation with preservation rates (Figure S14B, C).

We also calculated the preservation rates in Angara, Euramerica, Gondwana and South China paleoblocks. In Angara and South China, the preservation rates of macroplant are similar to the diversification trends, indicating that their diversity might be affected by preservation bias (Figure S15). In Euramerica and Gondwana, the preservation rates show different trends from their diversification.

**Extended drivers of diversification**

In different latitudinal zones, temperature ($G\mu$=-1.21, $\omega\mu$=0.99) and total area covered by sediment ($G\mu$=4.11, $\omega\mu$=0.97) are the most significant factors negative and positive correlation, respectively, with extinction in 0-30°N (Figure 4A and Table S5). $CO_2$, continental fragmentation, $O_2$, temperature and total area covered by sediment are the significant factors on origination

($\omega$>0.9) in 30-60°N (Figure 4A and Table S6). Dependence diversity, continental fragmentation and $O_2$ were significant correlation with origination ($\omega$>0.9) in 60-90°N, and continental fragmentation exhibit high negative correlation ($G\lambda$=-203.44) with origination and high positive correlation with ($G\mu$=50.60) with extinction (Table S7). In the 0-30°S, $CO_2$, continental fragmentation and $O_2$ are the significant drivers on extinction ($\omega$>0.9) (Table S8), similarly, continental fragmentation also shows high negative to origination ($G\lambda$=-30.27) and positive correlation with extinction ($G\mu$=133.91). $CO_2$ is the strong significant factor positively correlation with origination ($\omega\lambda$=0.91) in the 30-60°S (Table S9). In the 60-90°S, $CO_2$ level has a negative effect on origination ($\omega\lambda$=0.86), and $O_2$ also shows negative effect on extinction on extinction ($\omega\mu$=0.65) (Table S10).

In the different areas, In Angara, continental fragmentation shows strong significant factor negatively correlation with the origination ($G\lambda$=-89.83, $\omega\lambda$=0.93) and $O_2$ is the positive factors correlation with origination ($G\lambda$=7.19, $\omega\lambda$=0.93) (Table S11). In Euramerica, continental fragmentation shows high negatively to the extinction ($G\mu$=-30.51, $\omega\mu$=0.67) (Table S12). In Gondwana, continental fragmentation ($G\lambda$=17.29, $\omega\lambda$=0.66) and total area covered by sediment ($G\lambda$=3.45, $\omega\lambda$=0.98) are positive correlations with origination, and continental fragmentation shows positive correlation with extinction ($G\mu$=14.10, $\omega\mu$=0.63) (Table S13). In South China, continental fragmentation ($G\mu$=73.45, $\omega\mu$=0.95), $O_2$ ($G\mu$=23.19, $\omega\mu$=0.99) and total area covered by sediment ($G\mu$=10.86, $\omega\mu$=0.99) show high positive effect to origination.

**Dispersal in different latitudinal intervals**

We estimate the dispersal and extinction in different latitudinal intervals using dispersal-extinction-sampling (DES) models. Our results reveal variations in dispersal and extinction rates across different latitudinal intervals during the PTB. These results indicate that the Permo-Triassic interval does not exhibit a significant increase, thereby leading to marked shifts in species- and genus- diversity (figs S6 and S8).

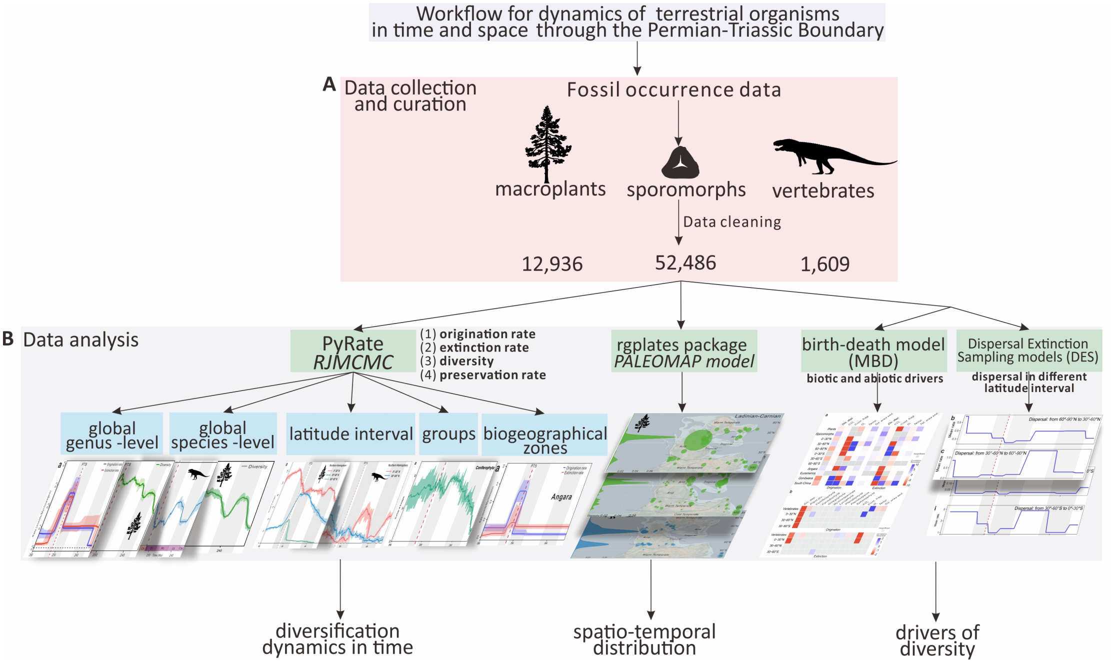


**Figure S1. Workflow for analyzing terrestrial biodiversity dynamics across the Permian–Triassic boundary.**

A. Data collection and curation. Fossil occurrence data for macroplants, sporomorphs and vertebrates were compiled and curated.

B. Data analysis. These datasets were analyzed using Bayesian farmwork to quantify origination, extinction, diversification and preservation rates, and reconstruct palaeogeographic distributions in R, investigate the biotic and abiotic drivers for diversity in multivariate birth-death (MBD) model and to calculate the dispersal in different latitudinal zones in dispersal extinction sampling models (DES).

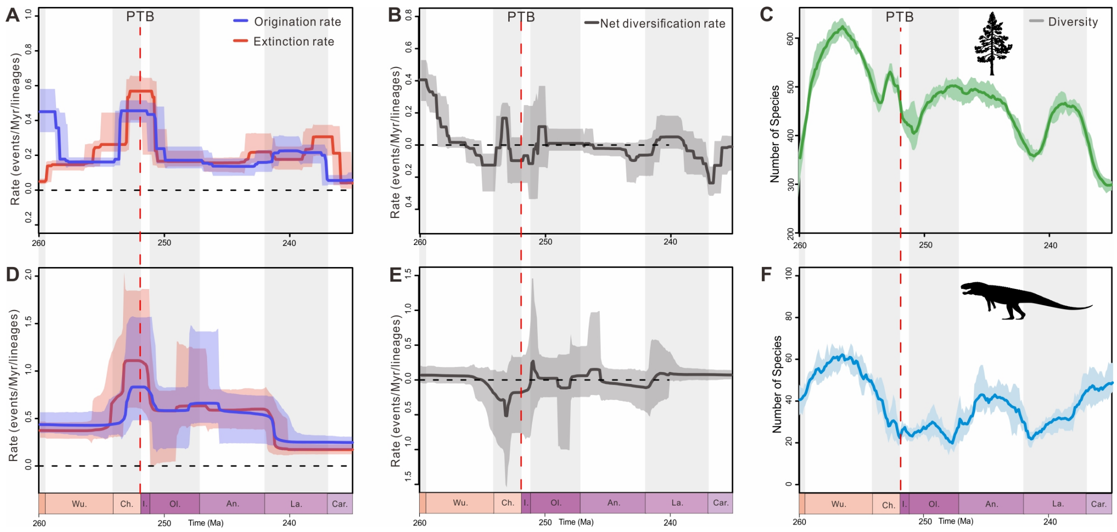


**Figure S2.** Estimated diversification and diversity dynamics of macroplant (A-C) and vertebrate (D-F) in species level. Solid lines indicate mean posterior density and the shaded areas show 95% highest posterior density (HPD) intervals. PTB: Permian-Triassic Boundary. Wo. Wordian, Cap. Capitanian, Wu. Wuchiapingian, Ch. Changhsingian, I. Induan, Ol. Olenekian, An. Anisian, La. Ladinian, Car. Carnian.

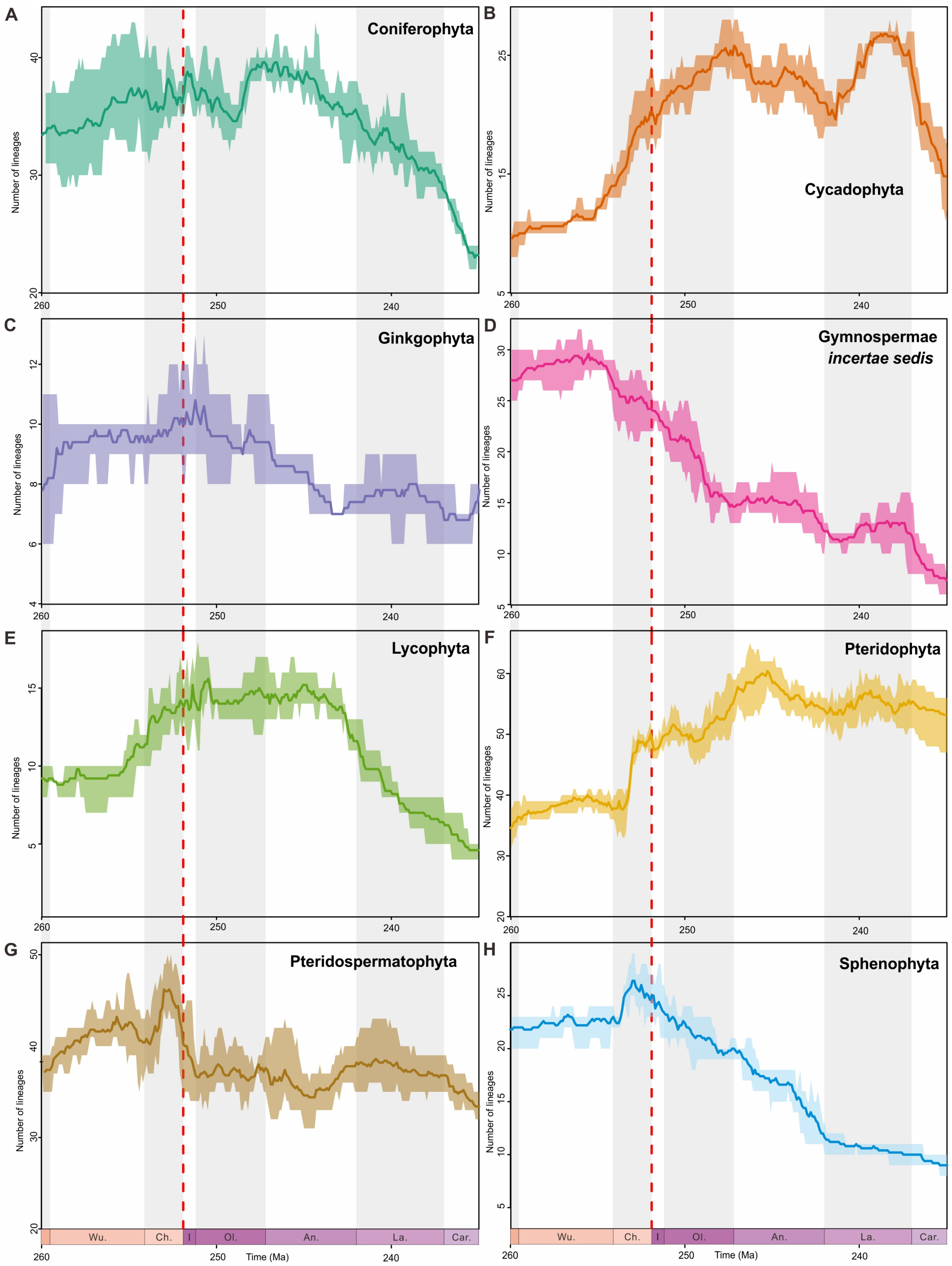


**Figure S3.** Diversity dynamics of main plant groups. PTB: Permian-Triassic Boundary. Wo. Wordian, Cap. Capitanian, Wu. Wuchiapingian, Ch. Changhsingian, I. Induan, Ol. Olenekian, An. Anisian, La. Ladinian, Car. Carnian.

$$\text{Origination }\% = \frac{N_{originated}}{N_{pre}} \times 100\%$$

$$\text{Extinction }\% = \frac{N_{extinct}}{N_{pre}} \times 100\%$$

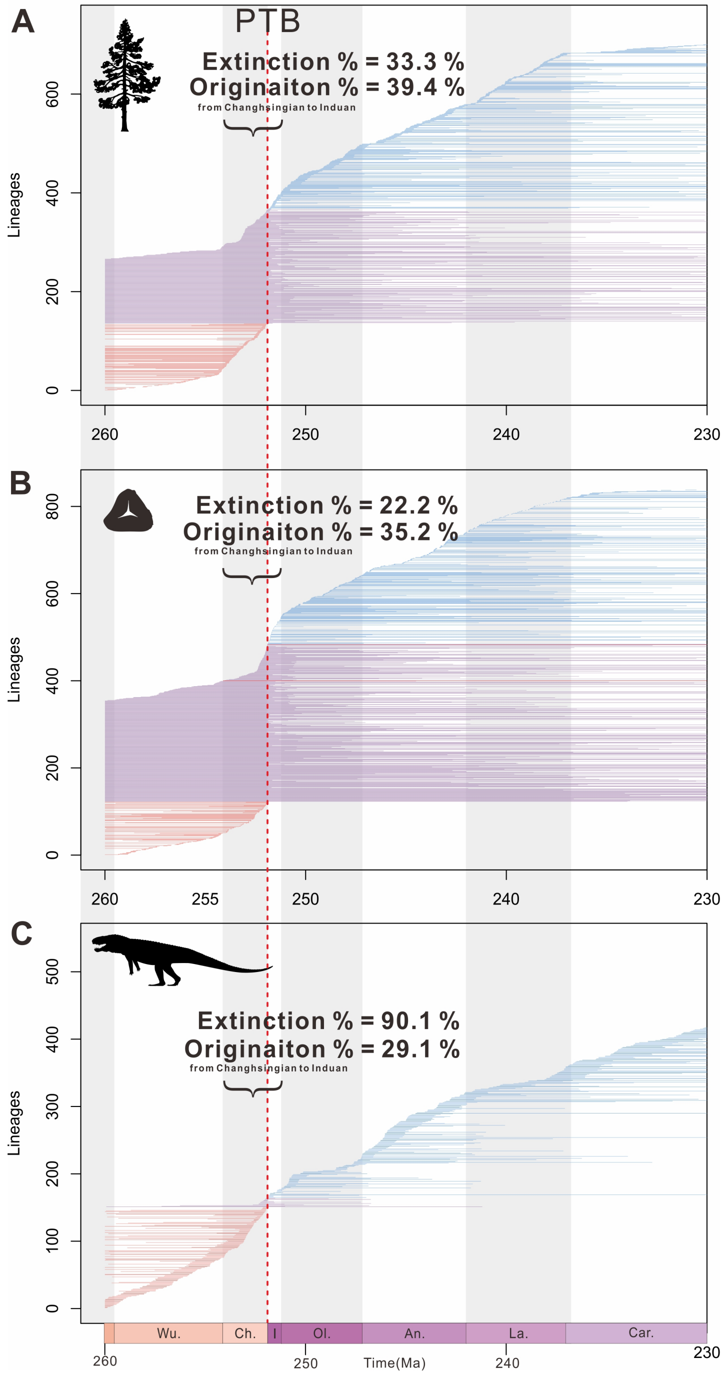


**Figure S4.** Genus's durations of macroplant, sporomorph and vertebrate estimated by PyRate simulations. In addition, extinction and origination percentages from Changhsingian to Induan are calculated based on genus's durations. PTB: Permian-Triassic Boundary.

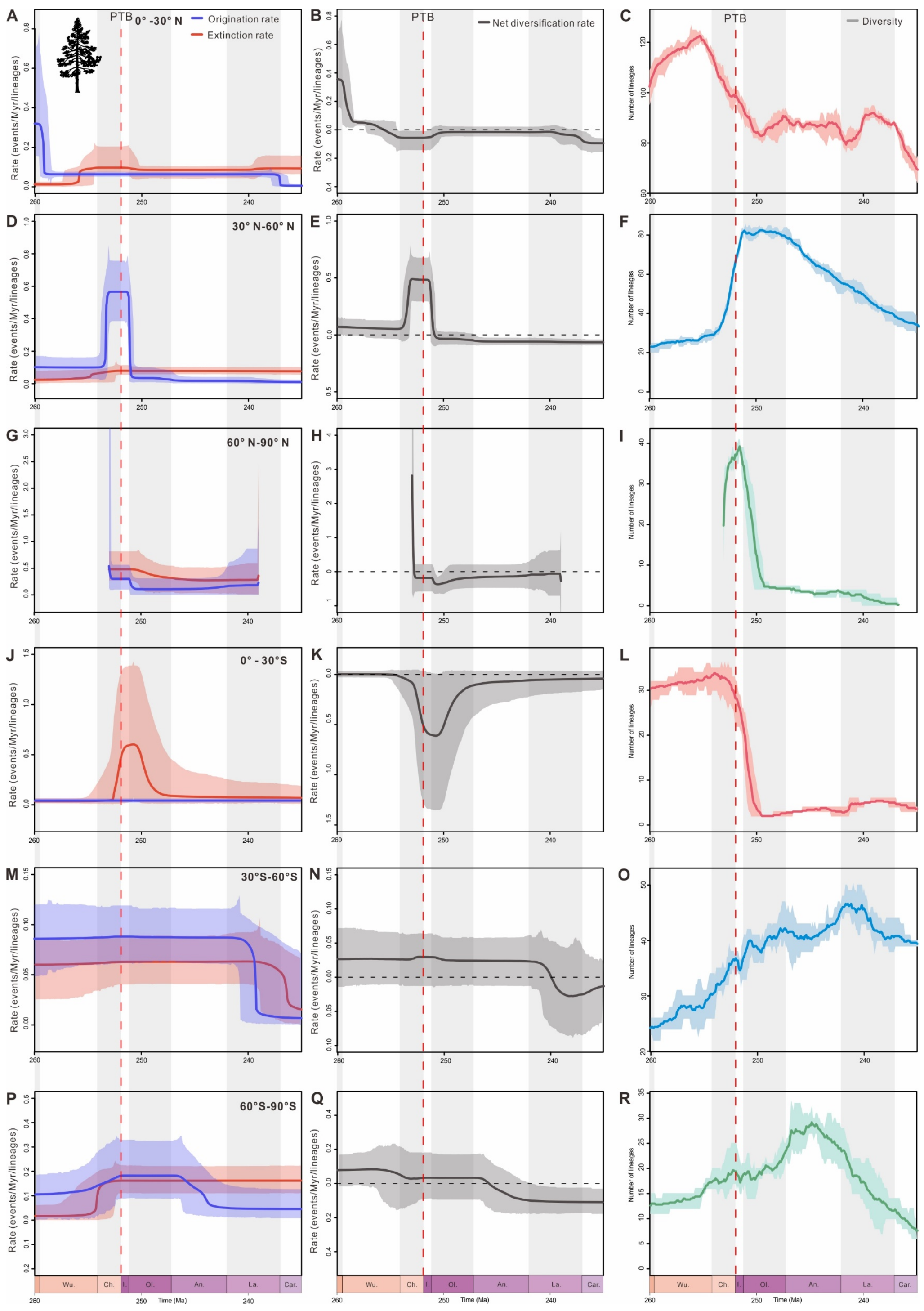


**Figure S5.** Estimated diversification and diversity dynamics of macroplant at different latitudinal zones. Solid lines indicate mean posterior density and the shaded areas show 95% highest posterior density (HPD) intervals. PTB: Permian-Triassic Boundary. Wo. Wordian, Cap. Capitanian, Wu. Wuchiapingian, Ch. Changhsingian, I. Induan, Ol. Olenekian, An. Anisian, La. Ladinian, Car. Carnian.

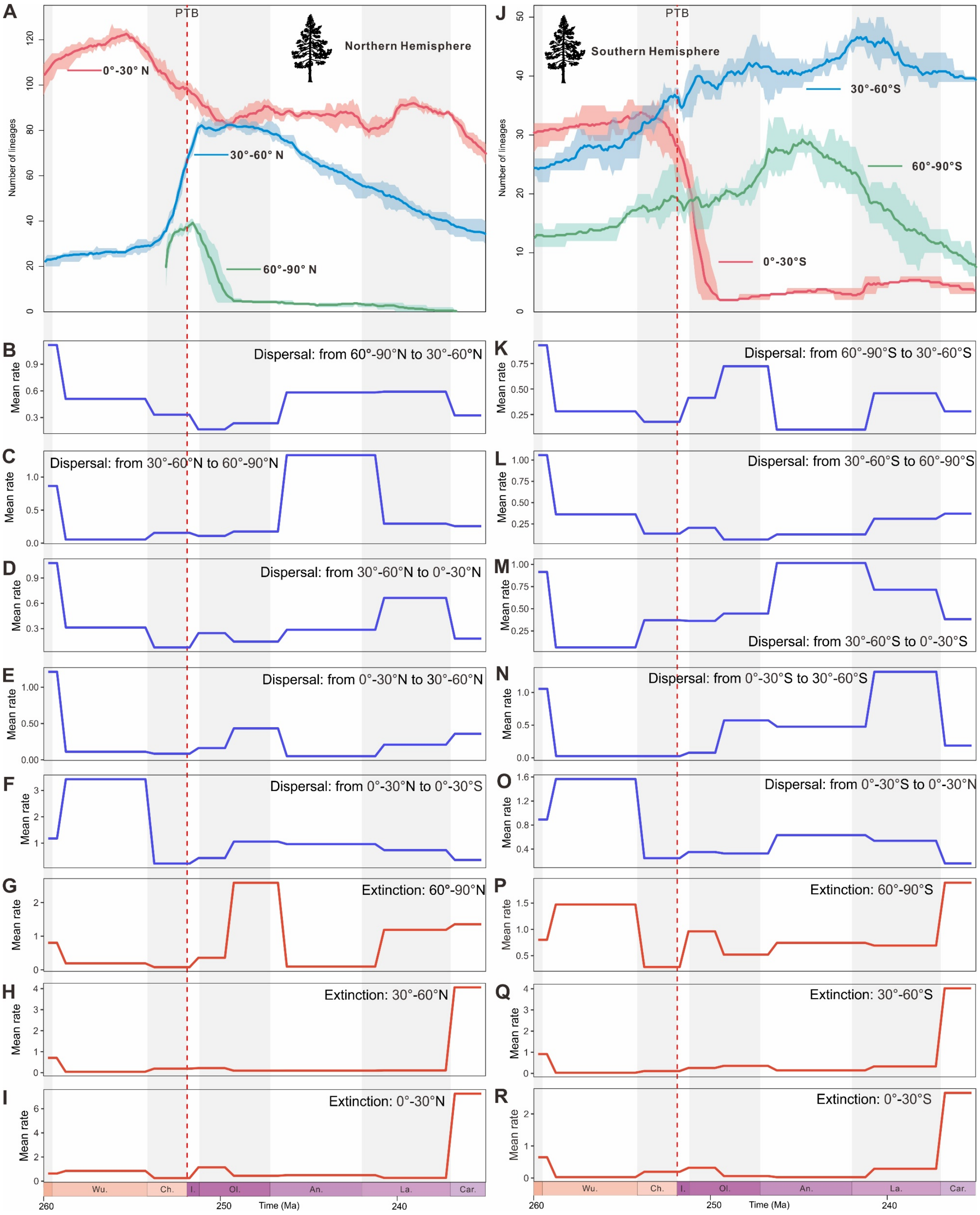


**Figure S6.** Plant diversity across different latitudinal intervals (A, J) are quantified based on palaeolatitudinal reconstructions. Dispersal and extinction rates are estimated among latitudinal bands, which show no significant changes across the Permian–Triassic boundary.

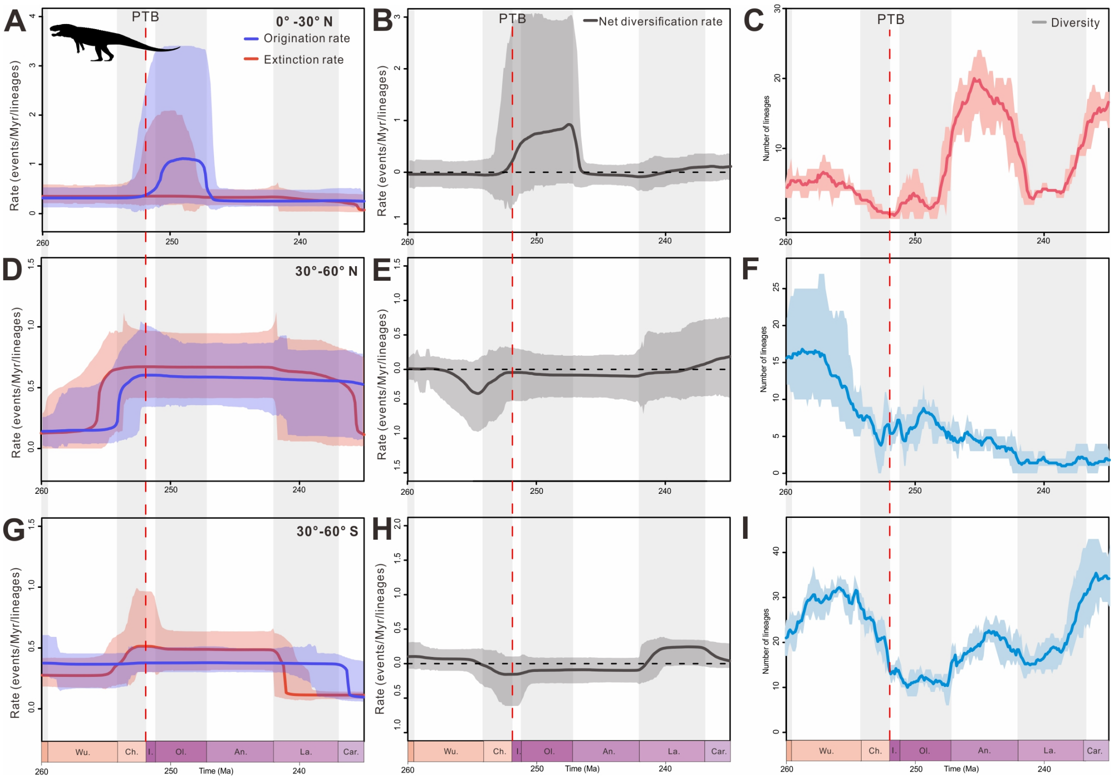


**Figure S7.** Estimated diversification and diversity dynamics of vertebrate at different latitudinal zones. Solid lines indicate mean posterior density and the shaded areas show 95% highest posterior density (HPD) intervals. PTB: Permian-Triassic Boundary. Wo. Wordian, Cap. Capitanian, Wu. Wuchiapingian, Ch. Changhsingian, I. Induan, Ol. Olenekian, An. Anisian, La. Ladinian, Car. Carnian.

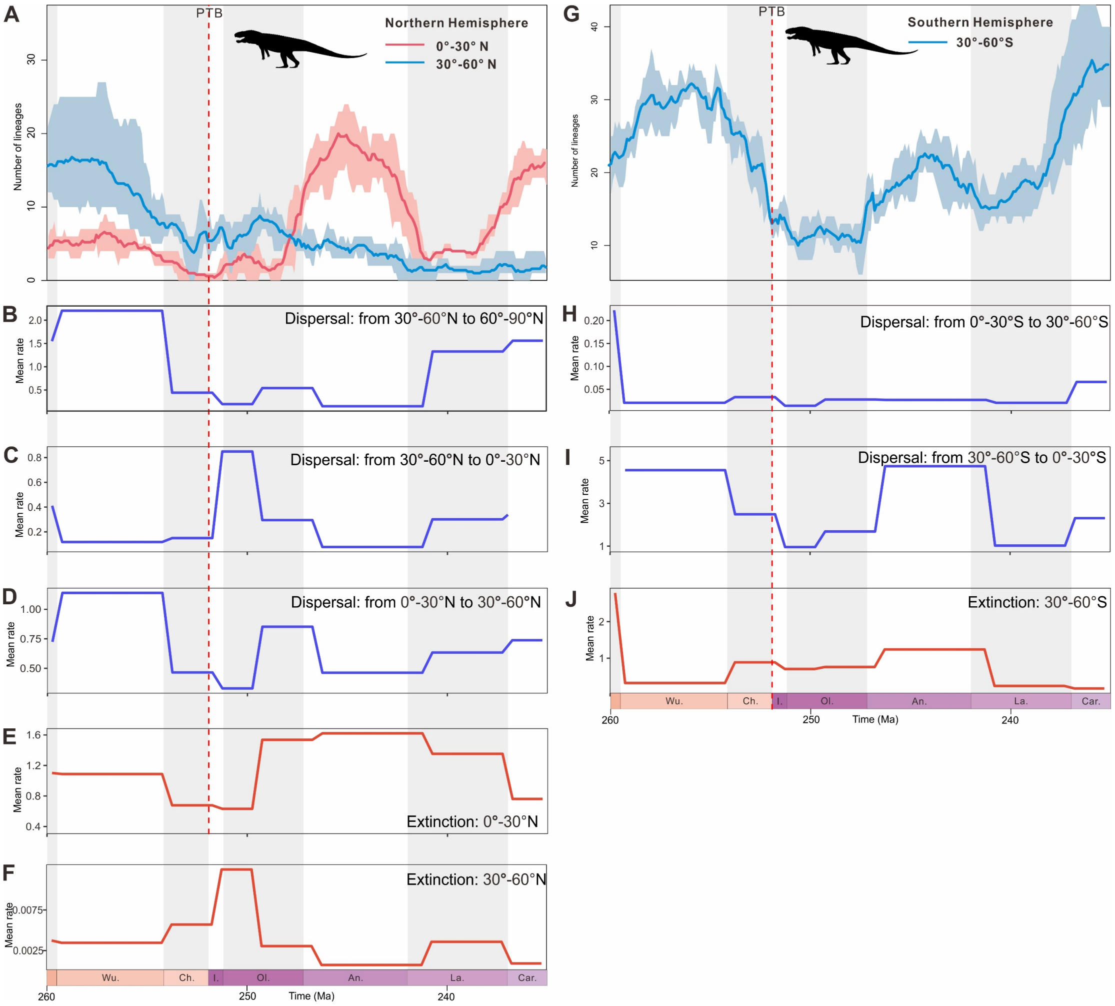


**Figure S8.** Vertebrate diversity across different latitudinal intervals (A, G) are quantified based on palaeolatitudinal reconstructions. Dispersal and extinction rates are estimated among latitudinal bands, which show no significant changes across the Permian–Triassic boundary, consistent with plant.

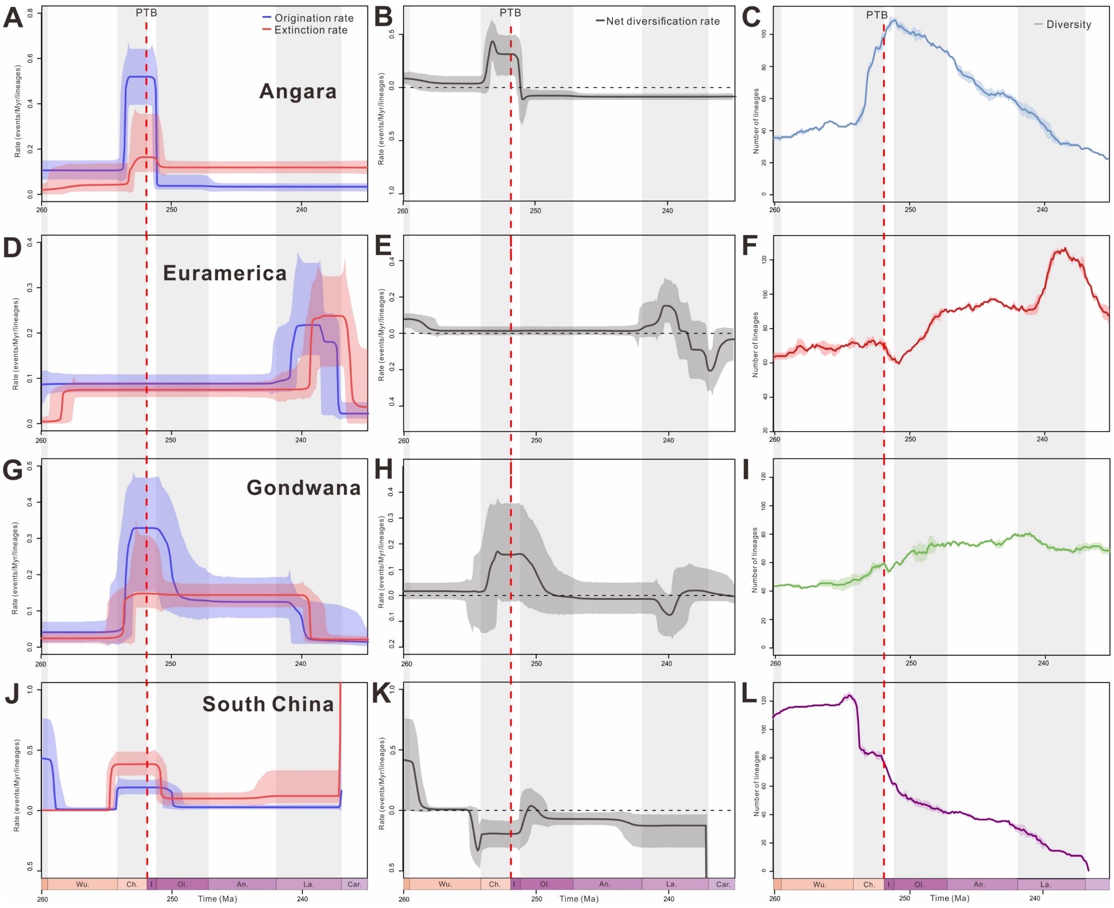


**Figure S9.** Estimated diversification and diversity dynamics of macroplant at Angara, Euramerica, Gondwana and South China zones. Solid lines indicate mean posterior density and the shaded areas show 95% highest posterior density (HPD) intervals. PTB: Permian-Triassic Boundary. Wo. Wordian, Cap. Capitanian, Wu. Wuchiapingian, Ch. Changhsingian, I. Induan, Ol. Olenekian, An. Anisian, La. Ladinian, Car. Carnian.

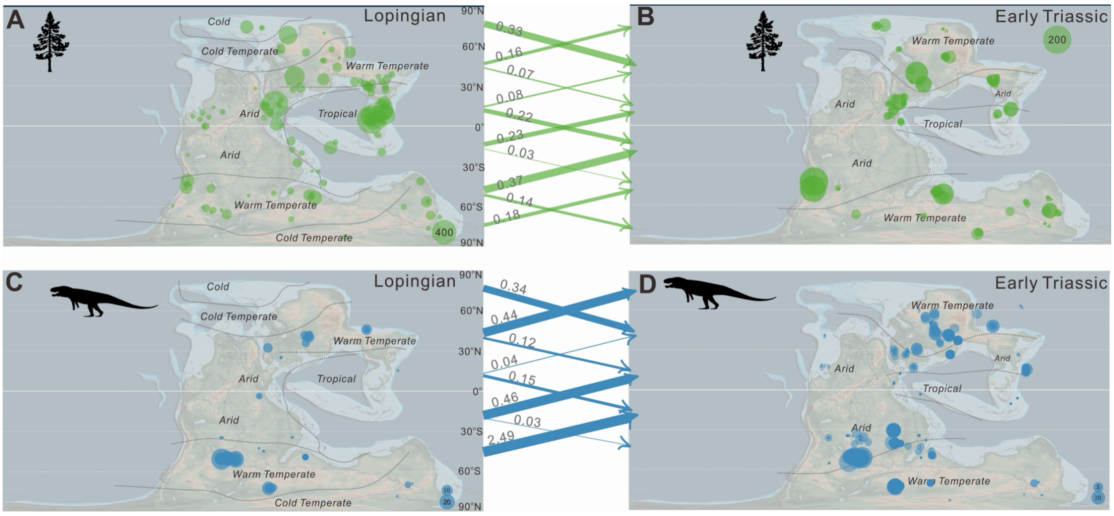


**Figure S10.** Dispersal rates among different latitudinal bands are shown as mean values at 251 Ma. The results indicate that dispersal among latitudinal zones in plants was relatively uniform, whereas vertebrates exhibit a more pronounced northward dispersal tendency. However, despite this northward shift, vertebrate dispersal did not fundamentally reshape overall diversity patterns.

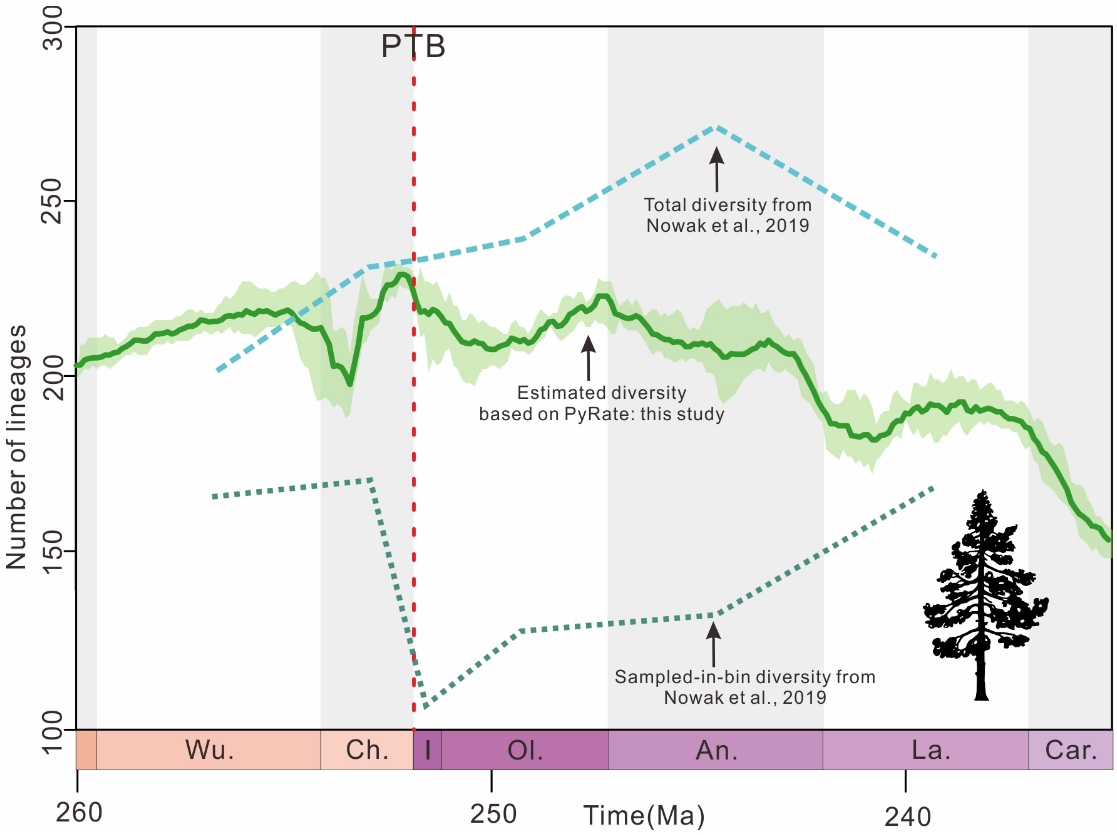


**Figure S11.** A comparison between the diversity patterns revealed in this study and those reported by Nowak et al. (2019) highlights differences in diversity estimates arising from the use of different methodological approaches.

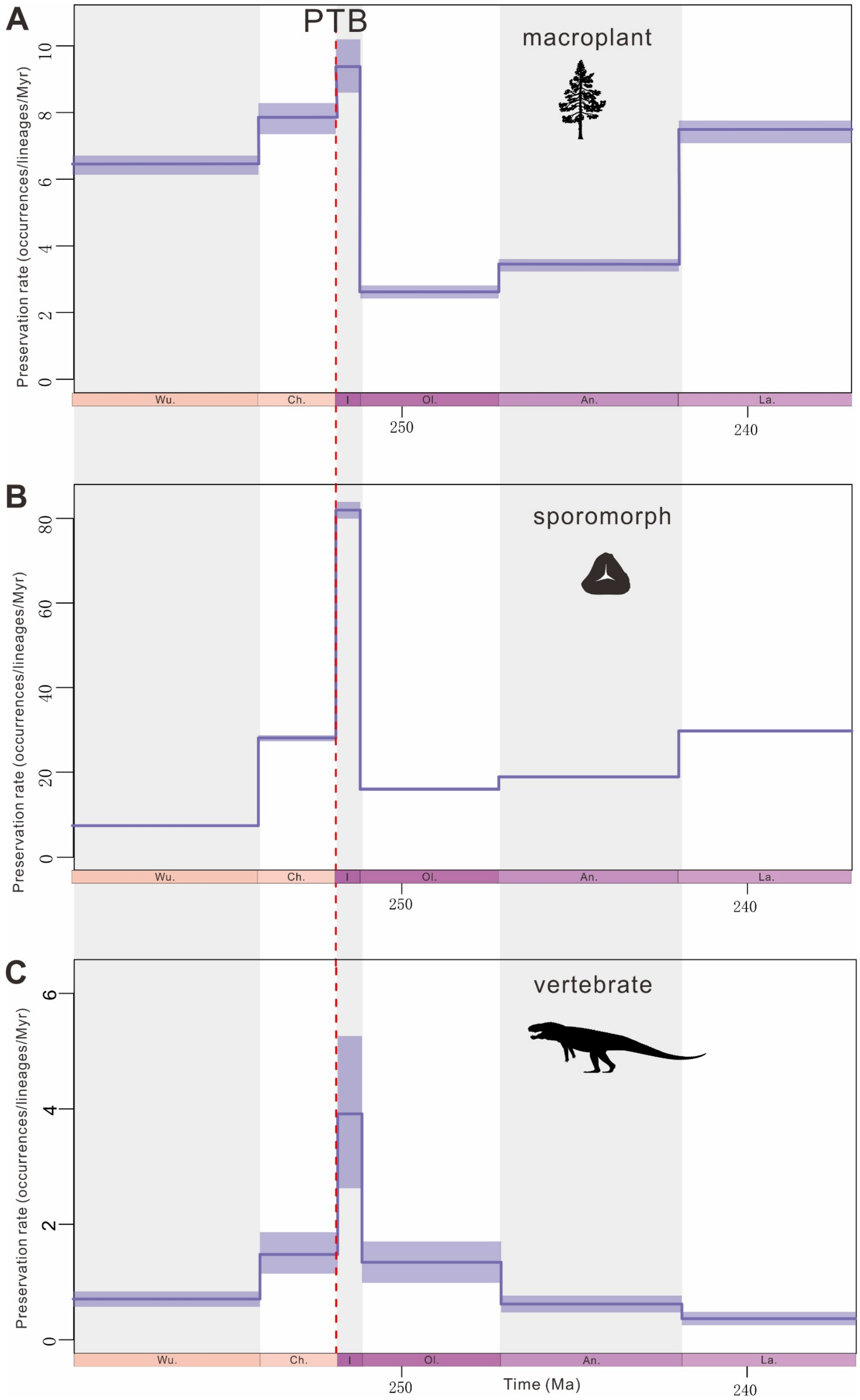


**Figure S12.** Temporal variation of the preservation rates of macroplant, sporomorph and vertebrate through the Permian-Triassic Boundary. The red dotted line indicates PTB. Wo. Wordian, Cap. Capitanian, Wu. Wuchiapingian, Ch. Changhsingian, I. Induan, Ol. Olenekian, An. Anisian, La. Ladinian, Car. Carnian.

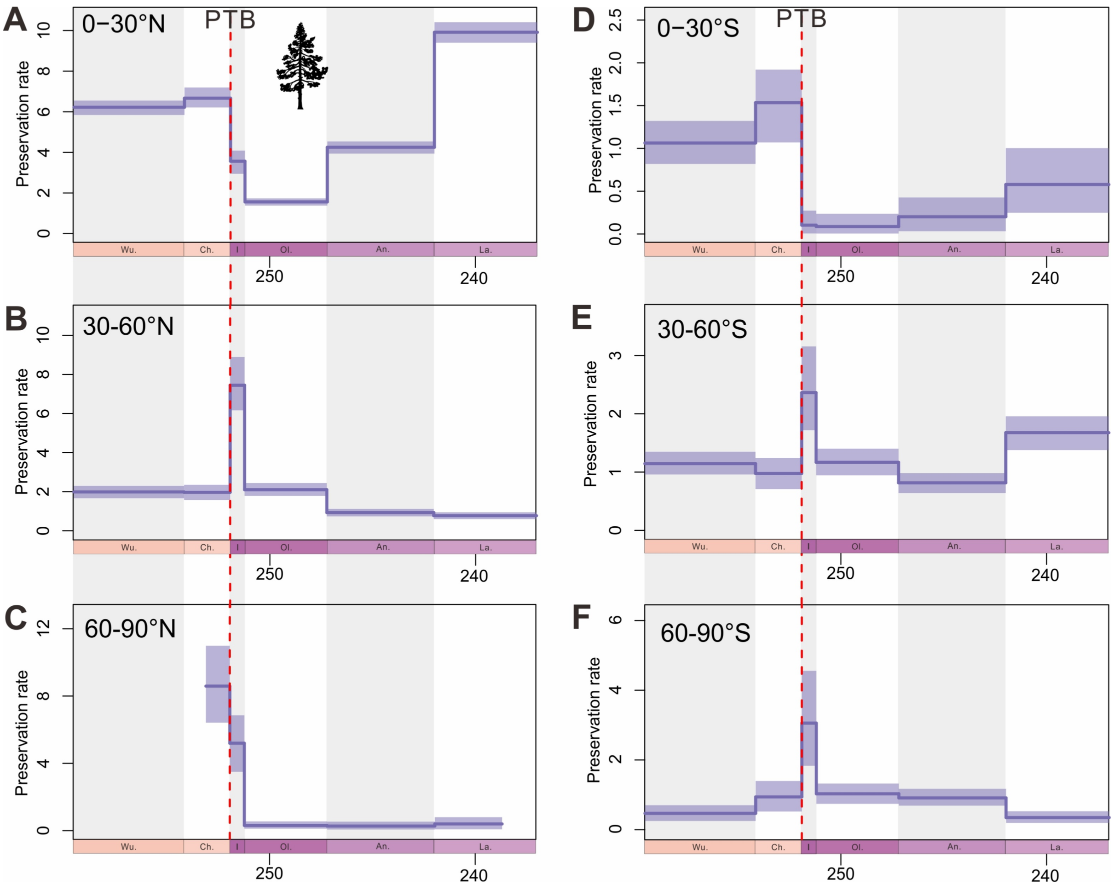


**Figure S13.** Temporal variation of the preservation rates of macroplant in different latitudinal zones. The red dotted line indicates PTB. Wo. Wordian, Cap. Capitanian, Wu. Wuchiapingian, Ch. Changhsingian, I. Induan, Ol. Olenekian, An. Anisian, La. Ladinian, Car. Carnian.

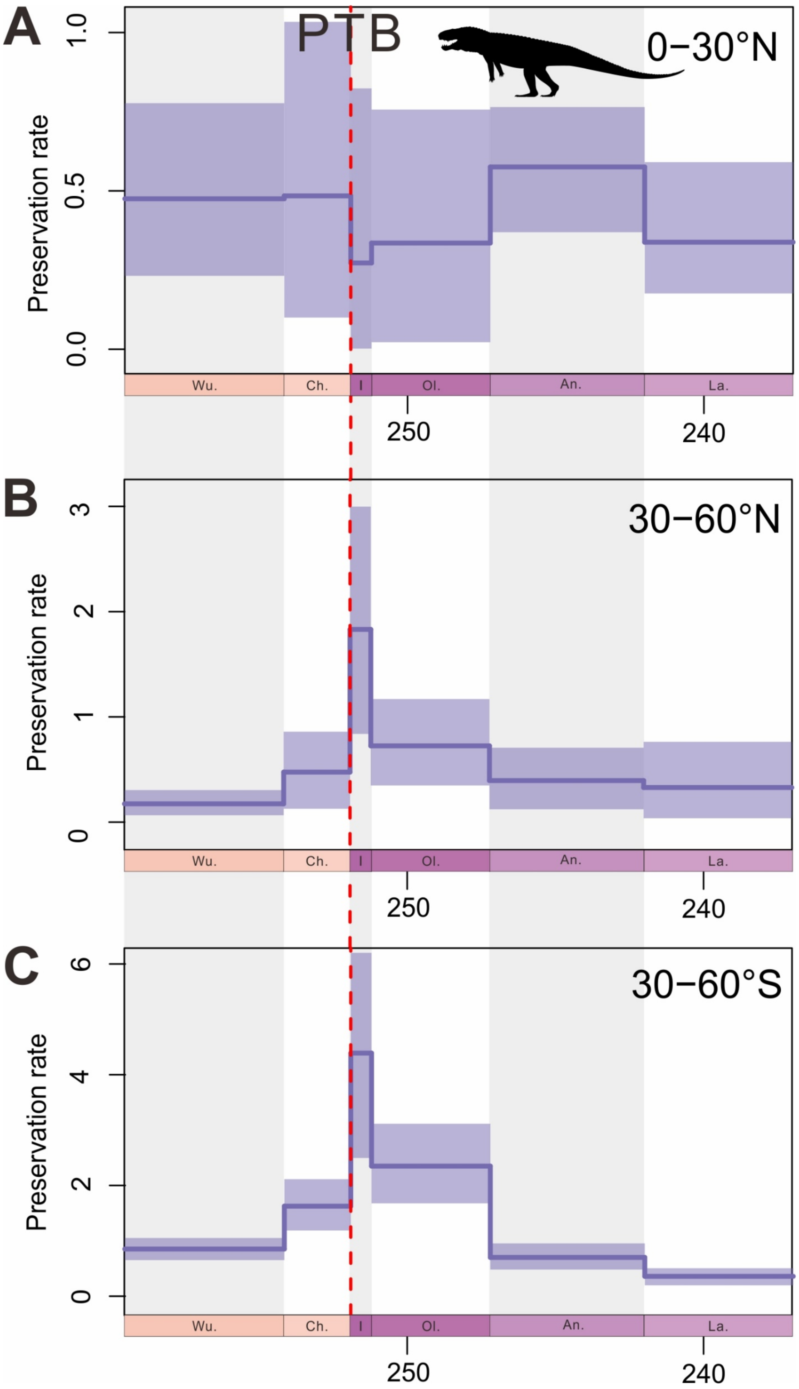


**Figure S14.** Temporal variation of the preservation rates of vertebrate in different latitudinal zones. The red dotted line indicates PTB. Wo. Wordian, Cap. Capitanian, Wu. Wuchiapingian, Ch. Changhsingian, I. Induan, Ol. Olenekian, An. Anisian, La. Ladinian, Car. Carnian.

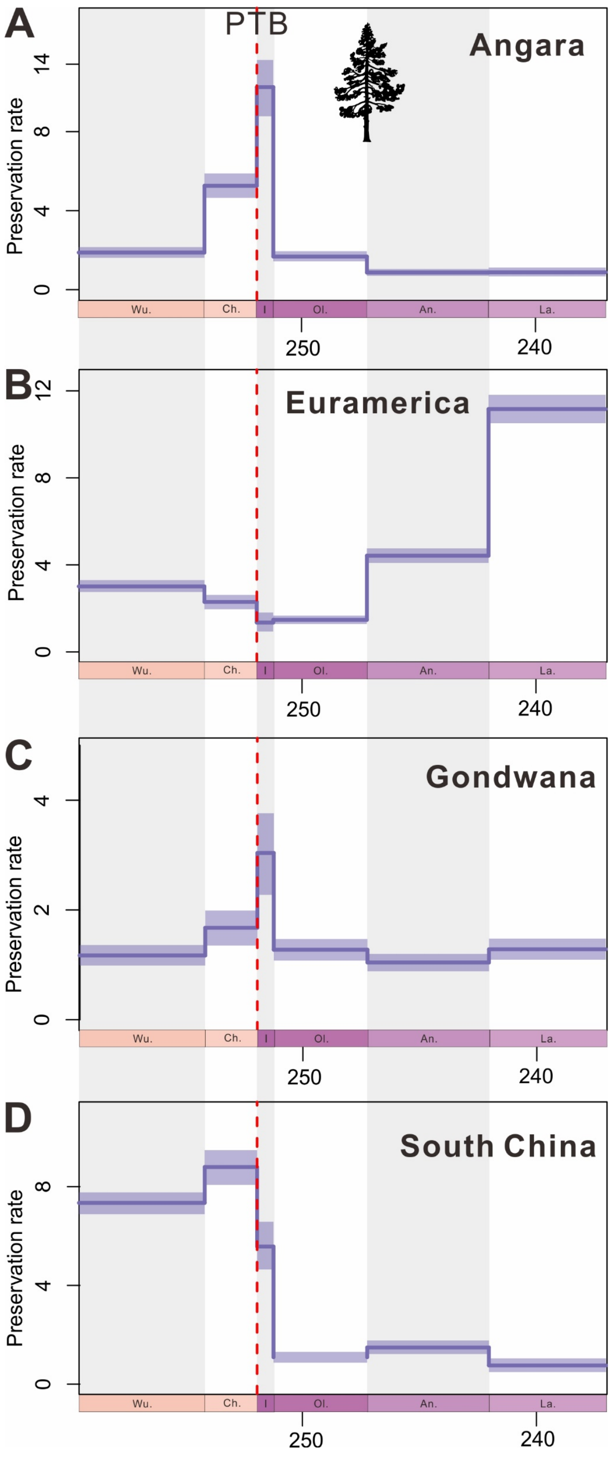


**Figure S15.** Temporal variation of the preservation rates of macroplant in Angara, Euramerica, Gondwana and South China areas. The red dotted line indicates PTB. Wo. Wordian, Cap. Capitanian, Wu. Wuchiapingian, Ch. Changhsingian, I. Induan, Ol. Olenekian, An. Anisian, La. Ladinian, Car. Carnian.

**Table S1.** Correlation parameters and Shrinkage weights of global plant using MBD model. Baseline origination and extinction rates (λ0 and μ0) and correlation parameters (Gλ and Gμ). The mean correlation parameters values were divided into pink (negative) and blue (positive). Shrinkage weights greater than 0.5 show significant evidence for correlation and are marked in orange. The strong support significant for bolded.

| Parameters | | | Median | Mean | CI_lower | CI_upper |
|---|---|---|---|---|---|---|
| Baseline rates | origination | λ0 | 0.000440279 | 0.001628646 | 5.27E-05 | 0.004209946 |
| | extinction | μ0 | 0.050582962 | 0.085378321 | 0.009681679 | 0.367776268 |
| Correlation parameters to origination | Diversity dependence | Gλ0_0 | -1.005188852 | -0.980700557 | -4.389807696 | 3.250958451 |
| | CO2 | Gλ0_1 | -0.000361907 | -0.000476034 | -0.001578202 | 0.00033769 |
| | O2 | Gλ0_3 | -3.290281964 | -3.226497356 | -5.910224878 | -0.170991165 |
| | Temperature | Gλ0_4 | -0.105345092 | -0.156661635 | -0.619686814 | 0.204977243 |
| | Total area covered by sediment | Gλ0_5 | 3.556501584 | 3.6254749 | 0.45563332 | 6.694561938 |
| Correlation parameters to extinction | Diversity dependence | Gμ0_0 | 2.337918867 | 2.567185847 | -0.188788782 | 6.29214378 |
| | CO2 | Gμ0_1 | -0.0002728 | -0.000320152 | -0.001047539 | 0.000198424 |
| | O2 | Gμ0_3 | 1.299559822 | 1.55481831 | -0.686663672 | 4.856843292 |
| | Temperature | Gμ0_4 | -0.850394912 | -0.864558296 | -1.32664134 | -0.482684318 |
| | Total area covered by sediment | Gμ0_5 | 3.497964728 | 3.249719689 | 0.094869203 | 5.567426727 |
| Shrinkage weights (origination) | Diversity dependence | ωλ0_0 | 0.841949346 | 0.71922037 | 0.01775821 | 0.996303181 |
| | CO2 | ωλ0_1 | 0.900813179 | 0.742421804 | 0.011333231 | 0.998074178 |
| | O2 | ωλ0_3 | 0.898104433 | 0.82402245 | 0.2 | 0.997219282 |
| | Temperature | ωλ0_4 | 0.916301424 | 0.766010638 | 0.019293415 | 0.998575441 |
| | Total area covered by sediment | ωλ0_5 | 0.990771844 | **0.96732229** | 0.759898347 | 0.999672073 |
| Shrinkage weights (extinction) | Diversity dependence | ωμ0_0 | 0.903463898 | 0.789842568 | 0.03983015 | 0.997668547 |
| | CO2 | ωμ0_1 | 0.83999912 | 0.698774119 | 0.007262761 | 0.996302883 |
| | O2 | ωμ0_3 | 0.767484292 | 0.648457186 | 0.004769266 | 0.995465293 |
| | Temperature | ωμ0_4 | 0.994423414 | **0.98918174** | 0.957469402 | 0.999791956 |
| | Total area covered by sediment | ωμ0_5 | 0.988841529 | **0.951672164** | 0.466185381 | 0.999656327 |

**Table S2.** Correlation parameters and Shrinkage weights of sporomorph using MBD model. Baseline origination and extinction rates (λ0 and μ0) and correlation parameters (Gλ and Gμ). The mean correlation parameters values were divided into pink (negative) and blue (positive). Shrinkage weights greater than 0.5 show significant evidence for correlation and are marked in orange. The strong support significant for bolded.

| Parameters | | | Median | Mean | CI_lower | CI_upper |
|---|---|---|---|---|---|---|
| Baseline rates | origination | λ0 | 0.000896685 | 0.002416224 | 0.000125327 | 0.004462747 |
| | extinction | μ0 | 0.307611684 | 0.393077386 | 0.065318216 | 1.259810871 |
| Correlation parameters to origination | Diversity dependence | Gλ0_0 | 3.83204011 | 3.522259715 | -0.012798015 | 5.68197199 |
| | CO2 | Gλ0_1 | -2.63E-05 | -0.000136511 | -0.001261291 | 0.000572396 |
| | O2 | Gλ0_3 | 0.008826868 | -0.254548263 | -3.573306668 | 1.598229565 |
| | Temperature | Gλ0_4 | 0.007310404 | 0.018176416 | -0.17538258 | 0.219781331 |
| | Total area covered by sediment | Gλ0_5 | 0.150670077 | 0.513102113 | -0.579799834 | 3.043290214 |
| Correlation parameters to extinction | Diversity dependence | Gμ0_0 | 6.061289663 | 5.931007112 | 2.698855668 | 8.938018498 |
| | CO2 | Gμ0_1 | -7.84E-06 | -0.0000312 | -0.001200115 | 0.001224681 |
| | O2 | Gμ0_3 | 3.288236321 | 3.591872723 | -0.10722681 | 8.540721215 |
| | Temperature | Gμ0_4 | -0.64105743 | -0.661844258 | -1.092125965 | - 0.271513822 |
| | Total area covered by sediment | Gμ0_5 | 0.023901505 | 0.131870906 | -1.154552243 | 1.824295206 |
| Shrinkage weights (origination) | Diversity dependence | ωλ0_0 | 0.930130398 | 0.870678434 | 0.258419194 | 0.99758962 |
| | CO2 | ωλ0_1 | 0.670280479 | 0.576226271 | 0.002410228 | 0.994975076 |
| | O2 | ωλ0_3 | 0.436632291 | 0.461540357 | 0.001082136 | 0.988817635 |
| | Temperature | ωλ0_4 | 0.594753048 | 0.543776369 | 0.002122384 | 0.993000331 |
| | Total area covered by sediment | ωλ0_5 | 0.703430883 | 0.598564903 | 0.003327441 | 0.996477723 |
| Shrinkage weights (extinction) | Diversity dependence | ωμ0_0 | 0.969455988 | **0.947869223** | 0.775504893 | 0.99889194 |
| | CO2 | ωμ0_1 | 0.802823829 | 0.646478083 | 0.003214829 | 0.997052261 |
| | O2 | ωμ0_3 | 0.882646531 | 0.781945425 | 0.056797943 | 0.996964239 |
| | Temperature | ωμ0_4 | 0.989912991 | **0.9800753** | 0.907782921 | 0.999637997 |
| | Total area covered by sediment | ωμ0_5 | 0.644444268 | 0.564622304 | 0.00230565 | 0.993377639 |

**Table S3.** Correlation parameters and Shrinkage weights of plant height in 0-6 m using MBD model. Baseline origination and extinction rates (λ0 and μ0) and correlation parameters (Gλ and Gμ). The mean correlation parameters values were divided into pink (negative) and blue (positive). Shrinkage weights greater than 0.5 show significant evidence for correlation and are marked in orange. The strong support significant for bolded.

| Parameters | | | Median | Mean | CI_lower | CI_upper |
|---|---|---|---|---|---|---|
| Baseline rates | origination | λ0 | 0.008135176 | 0.025000498 | 0.000191301 | 0.140647957 |
| | extinction | μ0 | 0.089008408 | 0.129187559 | 0.003894422 | 0.484614454 |
| Correlation parameters to origination | Diversity dependence | Gλ0_0 | 0.516128157 | 0.812400064 | -2.364855794 | 3.899294457 |
| | CO2 | Gλ0_1 | -0.001588159 | -0.001458761 | -0.0032272 | 0.000134145 |
| | O2 | Gλ0_3 | -0.18183284 | -0.456051186 | -4.083745038 | 2.002299097 |
| | Temperature | Gλ0_4 | -0.006241301 | -0.057535674 | -0.604530814 | 0.194181828 |
| | Total area covered by sediment | Gλ0_5 | 0.820410724 | 1.178601057 | -0.299204637 | 4.843371254 |
| Correlation parameters to extinction | Diversity dependence | Gμ0_0 | 3.558001923 | 3.283396451 | -0.674162007 | 7.966492487 |
| | CO2 | Gμ0_1 | -0.000565726 | -0.000634697 | -0.001819055 | 0.000117274 |
| | O2 | Gμ0_3 | 0.041798094 | 0.489120462 | -3.675879697 | 6.760098702 |
| | Temperature | Gμ0_4 | -0.245647977 | -0.312897783 | -1.034120236 | 0.033420764 |
| | Total area covered by sediment | Gμ0_5 | 0.011351863 | 0.276782225 | -1.97808512 | 4.226421931 |
| Shrinkage weights (origination) | Diversity dependence | ωλ0_0 | 0.689549539 | 0.585681781 | 0.002327761 | 0.991968919 |
| | CO2 | ωλ0_1 | 0.977492677 | 0.855581529 | 0.025555078 | 0.999378742 |
| | O2 | ωλ0_3 | 0.491266568 | 0.488279519 | 0.001775064 | 0.987707992 |
| | Temperature | ωλ0_4 | 0.661590858 | 0.580085769 | 0.002964904 | 0.996694761 |
| | Total area covered by sediment | ωλ0_5 | 0.898838386 | 0.735134493 | 0.008004877 | 0.998795209 |
| Shrinkage weights (extinction) | Diversity dependence | ωμ0_0 | 0.918911859 | 0.779496504 | 0.021207679 | 0.997305026 |
| | CO2 | ωμ0_1 | 0.907006523 | 0.740930932 | 0.010474301 | 0.997986566 |
| | O2 | ωμ0_3 | 0.678786131 | 0.58768319 | 0.004043128 | 0.9935423 |
| | Temperature | ωμ0_4 | 0.947464796 | 0.793553408 | 0.014809263 | 0.99914382 |
| | Total area covered by sediment | ωμ0_5 | 0.773763679 | 0.631770256 | 0.004185218 | 0.998236458 |

**Table S4.** Correlation parameters and Shrinkage weights of plant height more than 6 using MBD model. Baseline origination and extinction rates (λ0 and μ0) and correlation parameters (Gλ and Gμ). The mean correlation parameters values were divided into pink (negative) and blue (positive). Shrinkage weights greater than 0.5 show significant evidence for correlation and are marked in orange. The strong support significant for bolded.

| Parameters | | | Median | Mean | CI_lower | CI_upper |
|---|---|---|---|---|---|---|
| Baseline rates | origination | λ0 | 0.042977902 | 0.07651082 | 0.000784411 | 0.337286177 |
| | extinction | μ0 | 0.317418888 | 0.393935266 | 0.05449171 | 1.167448866 |
| Correlation parameters to origination | Diversity dependence | Gλ0_0 | 0.540866594 | 0.311378034 | -7.653698767 | 5.073085384 |
| | CO2 | Gλ0_1 | -0.002186832 | -0.002140526 | -0.003930693 | -3.41E-05 |
| | O2 | Gλ0_3 | -0.521648267 | -0.883215336 | -5.402001855 | 1.829424422 |
| | Temperature | Gλ0_4 | -0.034327205 | -0.146438045 | -0.910801979 | 0.126205888 |
| | Total area covered by sediment | Gλ0_5 | 0.201450749 | 1.232215958 | -1.388567303 | 7.349552395 |
| Correlation parameters to extinction | Diversity dependence | Gμ0_0 | 3.833802406 | 3.801923021 | 0.013839021 | 7.575420075 |
| | CO2 | Gμ0_1 | -0.000171078 | -0.000342634 | -0.00165284 | 0.000468117 |
| | O2 | Gμ0_3 | 0.901112556 | 1.806282747 | -1.985988378 | 7.656232932 |
| | Temperature | Gμ0_4 | -0.597264206 | -0.730118295 | -1.767284032 | -0.075211965 |
| | Total area covered by sediment | Gμ0_5 | 0.526725661 | 1.385636065 | -0.925003781 | 6.558714043 |
| Shrinkage weights (origination) | Diversity dependence | ωλ0_0 | 0.799796059 | 0.668206519 | 0.006190717 | 0.996562956 |
| | CO2 | ωλ0_1 | 0.990181243 | **0.951715923** | 0.367495721 | 0.999660696 |
| | O2 | ωλ0_3 | 0.6047925 | 0.547865066 | 0.001751434 | 0.994659489 |
| | Temperature | ωλ0_4 | 0.804818394 | 0.652361111 | 0.004233053 | 0.998789059 |
| | Total area covered by sediment | ωλ0_5 | 0.843300122 | 0.678273855 | 0.00413249 | 0.999411307 |
| Shrinkage weights (extinction) | Diversity dependence | ωμ0_0 | 0.941409633 | 0.877014262 | 0.268419547 | 0.998378415 |
| | CO2 | ωμ0_1 | 0.84166528 | 0.6823784 | 0.00500517 | 0.997513531 |
| | O2 | ωμ0_3 | 0.74517861 | 0.618241751 | 0.002791476 | 0.997071078 |
| | Temperature | ωμ0_4 | 0.990547994 | **0.961146125** | 0.704895217 | 0.999806845 |
| | Total area covered by sediment | ωμ0_5 | 0.91113067 | 0.720622167 | 0.006200417 | 0.999365439 |

**Table S5.** Correlation parameters and Shrinkage weights of plant in 0-30°N using MBD model. Baseline origination and extinction rates (λ0 and μ0) and correlation parameters (Gλ and Gμ). The mean correlation parameters values were divided into pink (negative) and blue (positive). Shrinkage weights greater than 0.5 show significant evidence for correlation and are marked in orange. The strong support significant for bolded.

| Parameters | | | Median | Mean | CI_lower | CI_upper |
|---|---|---|---|---|---|---|
| Baseline rates | origination | λ0 | 0.003926827 | 0.008087743 | 0.000204667 | 0.03637971 |
| | extinction | μ0 | 0.432923588 | 0.510516528 | 0.094788214 | 1.385832487 |
| Correlation parameters to origination | Diversity dependence | Gλ0_0 | -2.357219667 | -2.369853239 | -3.496462563 | -1.31486834 |
| | CO2 | Gλ0_1 | 2.67E-05 | 0.000108544 | -0.000926303 | 0.001468858 |
| | O2 | Gλ0_3 | 0.225646547 | 0.401109949 | -2.310492896 | 3.30661039 |
| | Temperature | Gλ0_4 | 0.159848905 | 0.171346083 | -0.151636471 | 0.534124294 |
| | Total area covered by sediment | Gλ0_5 | 1.028012833 | 1.114863722 | -0.446470033 | 3.215219789 |
| Correlation parameters to extinction | Diversity dependence | Gμ0_0 | 1.497896036 | 1.670054895 | -0.186381223 | 4.568316988 |
| | CO2 | Gμ0_1 | -0.000201629 | -0.000286371 | -0.00113031 | 0.000273647 |
| | O2 | Gμ0_3 | 2.859956668 | 2.897907972 | -0.352701218 | 6.556600612 |
| | Temperature | Gμ0_4 | -1.202006075 | -1.213494832 | -2.01849012 | -0.375901937 |
| | Total area covered by sediment | Gμ0_5 | 4.156934458 | 4.109916095 | 0.567158213 | 7.256797919 |
| Shrinkage weights (origination) | Diversity dependence | ωλ0_0 | 0.893076295 | 0.847349377 | 0.472500895 | 0.997089343 |
| | CO2 | ωλ0_1 | 0.785349937 | 0.651778175 | 0.0030333 | 0.996732729 |
| | O2 | ωλ0_3 | 0.618593399 | 0.553928272 | 0.003088365 | 0.992722696 |
| | Temperature | ωλ0_4 | 0.925974066 | 0.786282662 | 0.022041168 | 0.998447097 |
| | Total area covered by sediment | ωλ0_5 | 0.930041277 | 0.780625211 | 0.014579473 | 0.998559989 |
| Shrinkage weights (extinction) | Diversity dependence | ωμ0_0 | 0.817324347 | 0.70728353 | 0.017787456 | 0.996310743 |
| | CO2 | ωμ0_1 | 0.804303607 | 0.669330395 | 0.007920773 | 0.996866777 |
| | O2 | ωμ0_3 | 0.867649907 | 0.760782002 | 0.041385629 | 0.996886576 |
| | Temperature | ωμ0_4 | 0.996966815 | **0.991595791** | 0.958353563 | 0.99990651 |
| | Total area covered by sediment | ωμ0_5 | 0.992737695 | **0.968854612** | 0.780225811 | 0.999758109 |

**Table S6.** Correlation parameters and Shrinkage weights of plant in 30-60°N using MBD model. Baseline origination and extinction rates (λ0 and μ0) and correlation parameters (Gλ and Gμ). The mean correlation parameters values were divided into pink (negative) and blue (positive). Shrinkage weights greater than 0.5 show significant evidence for correlation and are marked in orange. The strong support significant for bolded.

| Parameters | | | Median | Mean | CI_lower | CI_upper |
|---|---|---|---|---|---|---|
| Baseline rates | origination | λ0 | 0.193618665 | 0.280166194 | 0.017666456 | 1.053506511 |
| | extinction | μ0 | 0.255219286 | 0.335211751 | 0.04340033 | 1.063259272 |
| Correlation parameters to origination | Diversity dependence | Gλ0_0 | 1.962929404 | 1.945133079 | 0.275445536 | 3.538869851 |
| | CO2 | Gλ0_1 | -0.004390141 | -0.004452481 | -0.007589819 | -0.001738484 |
| | O2 | Gλ0_3 | 13.53077819 | 13.47356634 | 4.466903528 | 23.45452538 |
| | Temperature | Gλ0_4 | -1.429473209 | -1.340021991 | -2.876043405 | 0.026242149 |
| | Total area covered by sediment | Gλ0_5 | 5.744187537 | 5.312378987 | -0.20276921 | 11.29414368 |
| Correlation parameters to extinction | Diversity dependence | Gμ0_0 | 1.138517213 | 1.224944233 | -0.204681867 | 3.257967127 |
| | CO2 | Gμ0_1 | -7.05E-06 | 7.21999E-05 | -0.001101818 | 0.001895263 |
| | O2 | Gμ0_3 | 0.44043638 | 1.288892399 | -2.695555514 | 8.829851356 |
| | Temperature | Gμ0_4 | -0.162193784 | -0.229970732 | -0.943920893 | 0.072235368 |
| | Total area covered by sediment | Gμ0_5 | -0.010122796 | 0.154937015 | -1.575527607 | 3.261878712 |
| Shrinkage weights (origination) | Diversity dependence | ωλ0_0 | 0.901038116 | 0.825170368 | 0.239195426 | 0.998457307 |
| | CO2 | ωλ0_1 | 0.997623929 | **0.994256062** | 0.976648933 | 0.999929198 |
| | O2 | ωλ0_3 | 0.989424589 | **0.974031792** | 0.846001495 | 0.999710292 |
| | Temperature | ωλ0_4 | 0.997705877 | **0.922039585** | 0.087537623 | 0.999939258 |
| | Total area covered by sediment | ωλ0_5 | 0.995756553 | **0.914393981** | 0.106673882 | 0.999900313 |
| Shrinkage weights (extinction) | Diversity dependence | ωμ0_0 | 0.833932365 | 0.71190875 | 0.017609262 | 0.997622368 |
| | CO2 | ωμ0_1 | 0.887291137 | 0.737016959 | 0.010258659 | 0.998620941 |
| | O2 | ωμ0_3 | 0.818998417 | 0.678954392 | 0.00746223 | 0.998304625 |
| | Temperature | ωμ0_4 | 0.949209188 | 0.819296512 | 0.028008043 | 0.999292261 |
| | Total area covered by sediment | ωμ0_5 | 0.876348823 | 0.727285507 | 0.010595926 | 0.999009268 |

**Table S7.** Correlation parameters and Shrinkage weights of plant in 60-90°N using MBD model. Baseline origination and extinction rates (λ0 and μ0) and correlation parameters (Gλ and Gμ). The mean correlation parameters values were divided into pink (negative) and blue (positive). Shrinkage weights greater than 0.5 show significant evidence for correlation and are marked in orange. The strong support significant for bolded.

| Parameters | | | Median | Mean | CI_lower | CI_upper |
|---|---|---|---|---|---|---|
| Baseline rates | origination | λ0 | 0.23023951 | 0.341236861 | 0.023651924 | 1.247659374 |
| | extinction | μ0 | 0.308798165 | 0.402372535 | 0.057767294 | 1.313466206 |
| Correlation parameters to origination | Diversity dependence | Gλ0_0 | -4.147918543 | -4.138047123 | -5.220333416 | -3.074487824 |
| | CO2 | Gλ0_1 | -8.71E-05 | -0.000994393 | -0.009068997 | 0.00163817 |
| | O2 | Gλ0_3 | 29.63673062 | 29.35223925 | 17.79810608 | 38.4060234 |
| | Temperature | Gλ0_4 | -0.0087739 | 0.001814955 | -0.376573062 | 0.545397659 |
| | Total area covered by sediment | Gλ0_5 | -0.316444761 | -0.594652204 | -2.753977497 | 0.886678207 |
| Correlation parameters to extinction | Diversity dependence | Gμ0_0 | 0.031403426 | 0.083742037 | -1.025871815 | 1.365732392 |
| | CO2 | Gμ0_1 | 2.91E-05 | 0.000197486 | -0.001027057 | 0.002562569 |
| | O2 | Gμ0_3 | 0.191556719 | 0.647924529 | -3.966585469 | 6.790988174 |
| | Temperature | Gμ0_4 | -0.009848657 | -0.016271251 | -0.382424172 | 0.353572713 |
| | Total area covered by sediment | Gμ0_5 | -0.048256553 | -0.19898377 | -2.044663096 | 1.172113946 |
| Shrinkage weights (origination) | Diversity dependence | ωλ0_0 | 0.946294969 | **0.924333079** | 0.729004953 | 0.998293144 |
| | CO2 | ωλ0_1 | 0.89125316 | 0.717664166 | 0.00677238 | 0.999775586 |
| | O2 | ωλ0_3 | 0.997166049 | **0.993644718** | 0.982086227 | 0.999910579 |
| | Temperature | ωλ0_4 | 0.803536953 | 0.668456478 | 0.0041327 | 0.998168679 |
| | Total area covered by sediment | ωλ0_5 | 0.835516094 | 0.682633838 | 0.006230581 | 0.998215104 |
| Shrinkage weights (extinction) | Diversity dependence | ωμ0_0 | 0.410511476 | 0.449784441 | 0.00151591 | 0.988668547 |
| | CO2 | ωμ0_1 | 0.795192961 | 0.657243505 | 0.003939959 | 0.998273803 |
| | O2 | ωμ0_3 | 0.689890141 | 0.593299486 | 0.004114169 | 0.995696383 |
| | Temperature | ωμ0_4 | 0.732070981 | 0.6197394 | 0.003192161 | 0.997967673 |
| | Total area covered by sediment | ωμ0_5 | 0.677285258 | 0.582246023 | 0.002407597 | 0.997500224 |

**Table S8.** Correlation parameters and Shrinkage weights of plant in 0-30°S using MBD model. Baseline origination and extinction rates (λ0 and μ0) and correlation parameters (Gλ and Gμ). The mean correlation parameters values were divided into pink (negative) and blue (positive). Shrinkage weights greater than 0.5 show significant evidence for correlation and are marked in orange. The strong support significant for bolded.

| Parameters | | | Median | Mean | CI_lower | CI_upper |
|---|---|---|---|---|---|---|
| Baseline rates | origination | λ0 | 0.208624585 | 0.296104112 | 0.036042125 | 1.089574482 |
| | extinction | μ0 | 0.292222046 | 0.370609587 | 0.029034466 | 1.146518668 |
| Correlation parameters to origination | Diversity dependence | Gλ0_0 | -0.279973749 | -0.457763903 | -2.480892061 | 1.078131214 |
| | CO2 | Gλ0_1 | -7.24E-05 | -0.000446754 | -0.003319609 | 0.00058294 |
| | O2 | Gλ0_3 | -0.095879104 | -0.173641957 | -4.090828616 | 4.025898401 |
| | Temperature | Gλ0_4 | -0.006951976 | -0.016177244 | -0.256706618 | 0.234958179 |
| | Total area covered by sediment | Gλ0_5 | -0.101732844 | -0.207947735 | -1.460563196 | 0.785040995 |
| Correlation parameters to extinction | Diversity dependence | Gμ0_0 | 0.370047839 | 0.61452485 | -1.27205923 | 3.079133482 |
| | CO2 | Gμ0_1 | -0.002134884 | -0.002104136 | -0.004537851 | 2.45E-05 |
| | O2 | Gμ0_3 | -8.462463008 | -8.261748972 | -15.45600583 | 0.217604345 |
| | Temperature | Gμ0_4 | -0.018551279 | -0.222741406 | -1.751449706 | 0.162311863 |
| | Total area covered by sediment | Gμ0_5 | 0.276256293 | 1.07554009 | -0.403546621 | 6.976850176 |
| Shrinkage weights (origination) | Diversity dependence | ωλ0_0 | 0.52087548 | 0.496658588 | 0.001438652 | 0.990501284 |
| | CO2 | ωλ0_1 | 0.782241881 | 0.634055025 | 0.003564913 | 0.998579108 |
| | O2 | ωλ0_3 | 0.582441212 | 0.531379869 | 0.001546654 | 0.992538659 |
| | Temperature | ωλ0_4 | 0.638382916 | 0.566987368 | 0.002507301 | 0.994834851 |
| | Total area covered by sediment | ωλ0_5 | 0.65280954 | 0.57092692 | 0.003173499 | 0.994008869 |
| Shrinkage weights (extinction) | Diversity dependence | ωμ0_0 | 0.590216958 | 0.538397629 | 0.001868981 | 0.992533189 |
| | CO2 | ωμ0_1 | 0.98879057 | **0.921733352** | 0.111667417 | 0.999689725 |
| | O2 | ωμ0_3 | 0.97075172 | **0.919435818** | 0.322002318 | 0.999217278 |
| | Temperature | ωμ0_4 | 0.741992189 | 0.625710846 | 0.00343159 | 0.999427888 |
| | Total area covered by sediment | ωμ0_5 | 0.761739347 | 0.636571076 | 0.003748782 | 0.99910461 |

**Table S9.** Correlation parameters and Shrinkage weights of plant in 30-60°S using MBD model. Baseline origination and extinction rates (λ0 and μ0) and correlation parameters (Gλ and Gμ). The mean correlation parameters values were divided into pink (negative) and blue (positive). Shrinkage weights greater than 0.5 show significant evidence for correlation and are marked in orange. The strong support significant for bolded.

| Parameters | | | Median | Mean | CI_lower | CI_upper |
|---|---|---|---|---|---|---|
| Baseline rates | origination | λ0 | 0.092542619 | 0.14037603 | 0.004230064 | 0.608232612 |
| | extinction | μ0 | 0.037031901 | 0.062383003 | 0.001429971 | 0.28920172 |
| Correlation parameters to origination | Diversity dependence | Gλ0_0 | -3.027861102 | -3.75996001 | -11.62256429 | 1.000553603 |
| | CO2 | Gλ0_1 | -0.001764591 | -0.001661281 | -0.003328141 | 3.78E-05 |
| | O2 | Gλ0_3 | -4.70666073 | -5.705987532 | -16.36357717 | 0.363428578 |
| | Temperature | Gλ0_4 | 0.003340426 | 0.02253014 | -0.385101598 | 0.503722401 |
| | Total area covered by sediment | Gλ0_5 | 1.504385997 | 1.828473486 | -0.128841336 | 5.398520678 |
| Correlation parameters to extinction | Diversity dependence | Gμ0_0 | 0.447328665 | 0.782104108 | -1.17273297 | 3.91815814 |
| | CO2 | Gμ0_1 | -0.000758358 | -0.000933542 | -0.002799544 | 0.00013626 |
| | O2 | Gμ0_3 | -0.736014062 | -1.276050847 | -5.713245204 | 1.243971627 |
| | Temperature | Gμ0_4 | 0.000934573 | 0.008684573 | -0.187459623 | 0.236573821 |
| | Total area covered by sediment | Gμ0_5 | 0.047006548 | 0.186077073 | -0.610503331 | 1.602203058 |
| Shrinkage weights (origination) | Diversity dependence | ωλ0_0 | 0.90571862 | 0.668713413 | 0.001281163 | 0.998511411 |
| | CO2 | ωλ0_1 | 0.983588441 | **0.912413769** | 0.148510524 | 0.999543245 |
| | O2 | ωλ0_3 | 0.923798966 | 0.715410292 | 0.002713059 | 0.998700952 |
| | Temperature | ωλ0_4 | 0.652686992 | 0.562755996 | 0.001411937 | 0.997366788 |
| | Total area covered by sediment | ωλ0_5 | 0.955017003 | 0.754298637 | 0.003182641 | 0.999330757 |
| Shrinkage weights (extinction) | Diversity dependence | ωμ0_0 | 0.58912278 | 0.530556572 | 0.001509369 | 0.991854463 |
| | CO2 | ωμ0_1 | 0.9375352 | 0.728047185 | 0.002066692 | 0.99905969 |
| | O2 | ωμ0_3 | 0.593116125 | 0.532129524 | 0.001174779 | 0.992735573 |
| | Temperature | ωμ0_4 | 0.532786544 | 0.501021972 | 0.000831543 | 0.993712834 |
| | Total area covered by sediment | ωμ0_5 | 0.531041942 | 0.505512521 | 0.000929718 | 0.993397645 |

**Table S10.** Correlation parameters and Shrinkage weights of plant in 60-90°S using MBD model. Baseline origination and extinction rates (λ0 and μ0) and correlation parameters (Gλ and Gμ). The mean correlation parameters values were divided into pink (negative) and blue (positive). Shrinkage weights greater than 0.5 show significant evidence for correlation and are marked in orange. The strong support significant for bolded.

<table>
<tr><th colspan="3">Parameters</th><th>Median</th><th>Mean</th><th>CI_lower</th><th>CI_upper</th></tr>
<tr><td rowspan="2">Baseline rates</td><td>origination</td><td>λ0</td><td>0.253385572</td><td>0.329116493</td><td>0.03735019</td><td>1.087605233</td></tr>
<tr><td>extinction</td><td>μ0</td><td>0.324235774</td><td>0.416334779</td><td>0.099662566</td><td>1.229942404</td></tr>
<tr><td rowspan="5">Correlation parameters to origination</td><td>Diversity dependence</td><td>Gλ0_0</td><td>0.315844172</td><td>0.496720024</td><td>-0.458774748</td><td>2.066962455</td></tr>
<tr><td>CO2</td><td>Gλ0_1</td><td>-0.002317571</td><td>-0.002186627</td><td>-0.004673117</td><td>3.13E-05</td></tr>
<tr><td>O2</td><td>Gλ0_3</td><td>-0.578254324</td><td>-0.979894168</td><td>-4.209959415</td><td>1.017736828</td></tr>
<tr><td>Temperature</td><td>Gλ0_4</td><td>-0.003456591</td><td>-0.03405628</td><td>-0.351308738</td><td>0.093442392</td></tr>
<tr><td>Total area covered by sediment</td><td>Gλ0_5</td><td>0.019010555</td><td>0.161521986</td><td>-0.408299178</td><td>1.561999536</td></tr>
<tr><td rowspan="5">Correlation parameters to extinction</td><td>Diversity dependence</td><td>Gμ0_0</td><td>0.00832747</td><td>0.038930953</td><td>-0.809774612</td><td>0.958041324</td></tr>
<tr><td>CO2</td><td>Gμ0_1</td><td>5.19E-07</td><td>1.35324E-05</td><td>-0.000685431</td><td>0.000795941</td></tr>
<tr><td>O2</td><td>Gμ0_3</td><td>-2.327323291</td><td>-2.288643265</td><td>-5.202980848</td><td>0.170458129</td></tr>
<tr><td>Temperature</td><td>Gμ0_4</td><td>-0.022183593</td><td>-0.067599826</td><td>-0.410314892</td><td>0.051609936</td></tr>
<tr><td>Total area covered by sediment</td><td>Gμ0_5</td><td>-0.008443111</td><td>0.026150135</td><td>-0.673361676</td><td>1.382451442</td></tr>
<tr><td rowspan="5">Shrinkage weights (origination)</td><td>Diversity dependence</td><td>ωλ0_0</td><td>0.304273634</td><td>0.370768343</td><td>0.000489886</td><td>0.968416301</td></tr>
<tr><td>CO2</td><td>ωλ0_1</td><td>0.988905341</td><td>0.862864164</td><td>0.004601947</td><td>0.999727708</td></tr>
<tr><td>O2</td><td>ωλ0_3</td><td>0.389259471</td><td>0.428664957</td><td>0.000554165</td><td>0.979624454</td></tr>
<tr><td>Temperature</td><td>ωλ0_4</td><td>0.307081054</td><td>0.395469121</td><td>0.000366269</td><td>0.989814774</td></tr>
<tr><td>Total area covered by sediment</td><td>ωλ0_5</td><td>0.260832034</td><td>0.367835748</td><td>0.000232303</td><td>0.984897107</td></tr>
<tr><td rowspan="5">Shrinkage weights (extinction)</td><td>Diversity dependence</td><td>ωμ0_0</td><td>0.140001537</td><td>0.264988649</td><td>0.000178566</td><td>0.951045612</td></tr>
<tr><td>CO2</td><td>ωμ0_1</td><td>0.306723022</td><td>0.400027902</td><td>0.000496248</td><td>0.988523872</td></tr>
<tr><td>O2</td><td>ωμ0_3</td><td>0.74254755</td><td>0.652090879</td><td>0.010925612</td><td>0.990468703</td></tr>
<tr><td>Temperature</td><td>ωμ0_4</td><td>0.449073302</td><td>0.464854439</td><td>0.000512174</td><td>0.992971679</td></tr>
<tr><td>Total area covered by sediment</td><td>ωμ0_5</td><td>0.282561846</td><td>0.370403413</td><td>0.000350526</td><td>0.981563666</td></tr>
</table>

**Table S11.** Correlation parameters and Shrinkage weights of plant in Angara using MBD model. Baseline origination and extinction rates (λ0 and μ0) and correlation parameters (Gλ and Gμ). The mean correlation parameters values were divided into pink (negative) and blue (positive). Shrinkage weights greater than 0.5 show significant evidence for correlation and are marked in orange. The strong support significant for bolded.

| Parameters | | | Median | Mean | CI_lower | CI_upper |
|---|---|---|---|---|---|---|
| Baseline rates | origination | λ0 | 0.351014 | 0.437923 | 0.061188 | 1.315388 |
| | extinction | μ0 | 0.186605 | 0.238018 | 0.043955 | 0.735305 |
| Correlation parameters to origination | Diversity dependence | Gλ0_0 | 1.606677 | 1.592157 | 0.440041 | 2.645355 |
| | CO2 | Gλ0_1 | -0.00366 | -0.0036 | -0.00556 | -0.00131 |
| | O2 | Gλ0_3 | 6.532055 | 7.189896 | 2.96589 | 13.85637 |
| | Temperature | Gλ0_4 | -0.19132 | -0.44469 | -1.73583 | 0.056913 |
| | Total area covered by sediment | Gλ0_5 | 0.400857 | 1.495262 | -0.58466 | 6.62973 |
| Correlation parameters to extinction | Diversity dependence | Gμ0_0 | 1.157281 | 1.162616 | 0.271271 | 2.05755 |
| | CO2 | Gμ0_1 | 1.94E-05 | 6.85E-05 | -0.00058 | 0.000901 |
| | O2 | Gμ0_3 | -0.42998 | -0.56309 | -2.54543 | 1.225194 |
| | Temperature | Gμ0_4 | -0.05763 | -0.07908 | -0.34322 | 0.084327 |
| | Total area covered by sediment | Gμ0_5 | -0.05145 | -0.08739 | -1.13528 | 1.195233 |
| Shrinkage weights (origination) | Diversity dependence | ωλ0_0 | 0.817813 | 0.755651 | 0.220668 | 0.994913 |
| | CO2 | ωλ0_1 | 0.996176 | **0.99052** | 0.955942 | 0.999879 |
| | O2 | ωλ0_3 | 0.96217 | **0.930698** | 0.683784 | 0.99891 |
| | Temperature | ωλ0_4 | 0.949857 | 0.769965 | 0.012416 | 0.999705 |
| | Total area covered by sediment | ωλ0_5 | 0.886059 | 0.703246 | 0.005809 | 0.999351 |
| Shrinkage weights (extinction) | Diversity dependence | ωμ0_0 | 0.746712 | 0.687078 | 0.123287 | 0.993321 |
| | CO2 | ωμ0_1 | 0.680959 | 0.588271 | 0.003749 | 0.994649 |
| | O2 | ωμ0_3 | 0.532902 | 0.512534 | 0.001884 | 0.990614 |
| | Temperature | ωμ0_4 | 0.779192 | 0.655325 | 0.006656 | 0.995883 |
| | Total area covered by sediment | ωμ0_5 | 0.675899 | 0.587572 | 0.00262 | 0.993784 |

**Table S12.** Correlation parameters and Shrinkage weights of plant in Euramerica using MBD model. Baseline origination and extinction rates (λ0 and μ0) and correlation parameters (Gλ and Gμ). The mean correlation parameters values were divided into pink (negative) and blue (positive). Shrinkage weights greater than 0.5 show significant evidence for correlation and are marked in orange. The strong support significant for bolded.

| Parameters | | | Median | Mean | CI_lower | CI_upper |
|---|---|---|---|---|---|---|
| Baseline rates | origination | λ0 | 0.011274 | 0.015985 | 0.001745 | 0.056472 |
| | extinction | μ0 | 0.37223 | 0.40916 | 0.141578 | 0.893377 |
| Correlation parameters to origination | Diversity dependence | Gλ0_0 | 0.814008 | 0.861352 | -0.2207 | 2.274436 |
| | CO2 | Gλ0_1 | -5.86E-05 | -0.00016 | -0.00093 | 0.000265 |
| | O2 | Gλ0_3 | -2.48921 | -2.54299 | -5.50924 | -0.01328 |
| | Temperature | Gλ0_4 | 0.070856 | 0.111381 | -0.05656 | 0.417056 |
| | Total area covered by sediment | Gλ0_5 | 0.190287 | 0.330154 | -0.436 | 1.443423 |
| Correlation parameters to extinction | Diversity dependence | Gμ0_0 | 0.316064 | 0.42408 | -0.36262 | 1.624737 |
| | CO2 | Gμ0_1 | -7.19E-07 | -8.13E-06 | -0.00042 | 0.000371 |
| | O2 | Gμ0_3 | 0.031401 | 0.239082 | -1.62424 | 3.241885 |
| | Temperature | Gμ0_4 | -0.15456 | -0.19147 | -0.70379 | 0.005939 |
| | Total area covered by sediment | Gμ0_5 | 0.011205 | 0.185403 | -0.74646 | 2.269541 |
| Shrinkage weights (origination) | Diversity dependence | ωλ0_0 | 0.529605 | 0.507902 | 0.003325 | 0.984192 |
| | CO2 | ωλ0_1 | 0.483103 | 0.481154 | 0.001114 | 0.988729 |
| | O2 | ωλ0_3 | 0.774952 | 0.689637 | 0.023176 | 0.992571 |
| | Temperature | ωλ0_4 | 0.735598 | 0.606797 | 0.00245 | 0.995804 |
| | Total area covered by sediment | ωλ0_5 | 0.587054 | 0.531972 | 0.001615 | 0.990962 |
| Shrinkage weights (extinction) | Diversity dependence | ωμ0_0 | 0.313222 | 0.377533 | 0.000792 | 0.967697 |
| | CO2 | ωμ0_1 | 0.323051 | 0.391089 | 0.000707 | 0.975429 |
| | O2 | ωμ0_3 | 0.282368 | 0.370925 | 0.00059 | 0.972487 |
| | Temperature | ωμ0_4 | 0.878099 | 0.768077 | 0.022773 | 0.997675 |
| | Total area covered by sediment | ωμ0_5 | 0.479022 | 0.483431 | 0.001238 | 0.991966 |

**Table S13.** Correlation parameters and Shrinkage weights of plant in Gondwana using MBD model. Baseline origination and extinction rates (λ0 and μ0) and correlation parameters (Gλ and Gμ). The mean correlation parameters values were divided into pink (negative) and blue (positive). Shrinkage weights greater than 0.5 show significant evidence for correlation and are marked in orange. The strong support significant for bolded.

| Parameters | | | Median | Mean | CI_lower | CI_upper |
|---|---|---|---|---|---|---|
| Baseline rates | origination | λ0 | 0.059109 | 0.100542 | 0.002946 | 0.427863 |
| | extinction | μ0 | 0.006389 | 0.015301 | 0.000362 | 0.083759 |
| Correlation parameters to origination | Diversity dependence | Gλ0_0 | -6.16621 | -6.14287 | -9.31232 | -2.80694 |
| | CO2 | Gλ0_1 | -0.00157 | -0.00153 | -0.00262 | -0.00022 |
| | O2 | Gλ0_3 | -11.3892 | -11.3824 | -15.7133 | -6.99256 |
| | Temperature | Gλ0_4 | -0.00496 | -0.01972 | -0.4269 | 0.341662 |
| | Total area covered by sediment | Gλ0_5 | 3.405179 | 3.453792 | 1.821991 | 5.291678 |
| Correlation parameters to extinction | Diversity dependence | Gμ0_0 | 0.001811 | -0.01873 | -2.82605 | 2.770662 |
| | CO2 | Gμ0_1 | -0.00088 | -0.00086 | -0.00186 | 5.83E-05 |
| | O2 | Gμ0_3 | -4.44107 | -4.39748 | -8.08336 | -0.44188 |
| | Temperature | Gμ0_4 | -0.02861 | -0.07896 | -0.59549 | 0.245386 |
| | Total area covered by sediment | Gμ0_5 | 1.780019 | 1.901801 | 0.236776 | 4.28022 |
| Shrinkage weights (origination) | Diversity dependence | ωλ0_0 | 0.976263 | **0.957952** | 0.815123 | 0.99929 |
| | CO2 | ωλ0_1 | 0.984203 | **0.958539** | 0.738332 | 0.999548 |
| | O2 | ωλ0_3 | 0.98533 | **0.976617** | 0.90599 | 0.999513 |
| | Temperature | ωλ0_4 | 0.86131 | 0.712609 | 0.007884 | 0.998035 |
| | Total area covered by sediment | ωλ0_5 | 0.990743 | **0.98412** | 0.932526 | 0.999669 |
| Shrinkage weights (extinction) | Diversity dependence | ωμ0_0 | 0.74149 | 0.629502 | 0.005309 | 0.995992 |
| | CO2 | ωμ0_1 | 0.961409 | 0.885111 | 0.161064 | 0.99894 |
| | O2 | ωμ0_3 | 0.940124 | 0.884888 | 0.394055 | 0.998438 |
| | Temperature | ωμ0_4 | 0.884916 | 0.736412 | 0.010854 | 0.998272 |
| | Total area covered by sediment | ωμ0_5 | 0.974311 | **0.938726** | 0.633185 | 0.999263 |

**Table S14.** Correlation parameters and Shrinkage weights of plant in South China using MBD model. Baseline origination and extinction rates (λ0 and μ0) and correlation parameters (Gλ and Gμ). The mean correlation parameters values were divided into pink (negative) and blue (positive). Shrinkage weights greater than 0.5 show significant evidence for correlation and are marked in orange. The strong support significant for bolded.

| Parameters | | | Median | Mean | CI_lower | CI_upper |
|---|---|---|---|---|---|---|
| Baseline rates | origination | λ0 | 0.557849 | 0.691749 | 0.072907 | 2.135001 |
| | extinction | μ0 | 0.957828 | 1.118911 | 0.225064 | 2.910108 |
| Correlation parameters to origination | Diversity dependence | Gλ0_0 | -3.98514 | -3.98986 | -4.82477 | -3.1573 |
| | CO2 | Gλ0_1 | 0.000525 | 0.000743 | -0.00097 | 0.003121 |
| | O2 | Gλ0_3 | 23.27112 | 23.19889 | 17.43426 | 28.77426 |
| | Temperature | Gλ0_4 | -3.00753 | -2.99729 | -3.90113 | -2.04386 |
| | Total area covered by sediment | Gλ0_5 | 10.89998 | 10.86365 | 7.081067 | 14.31435 |
| Correlation parameters to extinction | Diversity dependence | Gμ0_0 | 0.617441 | 0.689366 | -0.80539 | 2.494428 |
| | CO2 | Gμ0_1 | 0.000619 | 0.00075 | -0.00062 | 0.002665 |
| | O2 | Gμ0_3 | 8.219605 | 8.239754 | 2.815001 | 13.76914 |
| | Temperature | Gμ0_4 | -3.01821 | -3.029 | -3.90256 | -2.22133 |
| | Total area covered by sediment | Gμ0_5 | 12.13097 | 12.13254 | 8.907587 | 15.49278 |
| Shrinkage weights (origination) | Diversity dependence | ωλ0_0 | 0.979165 | **0.958283** | 0.799043 | 0.99966 |
| | CO2 | ωλ0_1 | 0.978698 | 0.879098 | 0.08047 | 0.999715 |
| | O2 | ωλ0_3 | 0.996928 | **0.994984** | 0.979855 | 0.999896 |
| | Temperature | ωλ0_4 | 0.999565 | **0.999113** | 0.997165 | 0.999983 |
| | Total area covered by sediment | ωλ0_5 | 0.99905 | **0.998042** | 0.993527 | 0.999966 |
| Shrinkage weights (extinction) | Diversity dependence | ωμ0_0 | 0.873783 | 0.723405 | 0.010757 | 0.999126 |
| | CO2 | ωμ0_1 | 0.97829 | 0.879135 | 0.079667 | 0.999668 |
| | O2 | ωμ0_3 | 0.987177 | **0.968443** | 0.833976 | 0.999715 |
| | Temperature | ωμ0_4 | 0.999559 | **0.998876** | 0.997278 | 0.999986 |
| | Total area covered by sediment | ωμ0_5 | 0.999204 | **0.998429** | 0.995219 | 0.999974 |

**Table S15.** Correlation parameters and Shrinkage weights of global vertebrate using MBD model. Baseline origination and extinction rates (λ0 and μ0) and correlation parameters (Gλ and Gμ). The mean correlation parameters values were divided into pink (negative) and blue (positive). Shrinkage weights greater than 0.5 show significant evidence for correlation and are marked in orange. The strong support significant for bolded.

| Parameters | | | Median | Mean | CI_lower | CI_upper |
|---|---|---|---|---|---|---|
| Baseline rates | origination | λ0 | 0.351253216 | 0.455827863 | 0.057087267 | 1.41900618 |
| | extinction | μ0 | 0.606952715 | 0.694193986 | 0.148074544 | 1.728314008 |
| Correlation parameters to origination | Diversity dependence | Gλ0_0 | -2.147834015 | -2.11254121 | -3.24419723 | -0.861588563 |
| | Plant diversity | Gλ0_1 | 0.022573608 | 0.03169492 | -0.016368119 | 0.115804486 |
| | Coniferophyta | Gλ0_2 | -0.006472777 | -0.01644048 | -0.105193184 | 0.045554058 |
| | Cycadophyta | Gλ0_3 | -0.011426394 | -0.027049708 | -0.176684909 | 0.069015733 |
| | Ginkgophyta | Gλ0_4 | -0.000300702 | -0.00314703 | -0.090192638 | 0.064313084 |
| | Gymnospermae | Gλ0_5 | 0.002307545 | 0.004161894 | -0.077591416 | 0.080610245 |
| | Lycophyta | Gλ0_6 | 0.059753863 | 0.057417981 | -0.003540999 | 0.128586005 |
| | Pteridophyta | Gλ0_7 | 0.000621004 | 0.004240878 | -0.036626417 | 0.062254275 |
| | Pteridospermatophyta | Gλ0_8 | 0.00387721 | 0.01037909 | -0.070130357 | 0.109822174 |
| | Sphenophyta | Gλ0_9 | 0.00536403 | 0.013963373 | -0.067143007 | 0.113432529 |
| | CO2 | Gλ0_10 | 4.66E-06 | 0.0000388 | -0.000431579 | 0.000693797 |
| | O2 | Gλ0_12 | 6.091363811 | 5.786319418 | -0.206683604 | 12.87014235 |
| | Temp. | Gλ0_13 | -0.299458289 | -0.349753075 | -0.998115453 | 0.0225192 |
| | Total area covered by sed. | Gλ0_14 | -0.011744145 | -0.045832025 | -1.768122045 | 1.712375177 |
| Correlation parameters to extinction | Diversity dependence | Gμ0_0 | -1.551888888 | -1.543345733 | -2.428758196 | -0.587676897 |
| | Plant diversity | Gμ0_1 | 0.004043555 | 0.017657879 | -0.030613484 | 0.142232822 |
| | Coniferophyta | Gμ0_2 | -0.061113289 | -0.069765017 | -0.189702807 | 0.005270645 |
| | Cycadophyta | Gμ0_3 | -0.055521934 | -0.071615087 | -0.26214966 | 0.067259854 |
| | Ginkgophyta | Gμ0_4 | -0.007726001 | -0.028320889 | -0.186785367 | 0.039460608 |
| | Gymnospermae | Gμ0_5 | 0.01005833 | 0.022898139 | -0.066156343 | 0.144267368 |
| | Lycophyta | Gμ0_6 | -0.00344349 | -0.010139629 | -0.071993998 | 0.022206449 |
| | Pteridophyta | Gμ0_7 | 0.010312033 | 0.017624785 | -0.014909522 | 0.079124095 |
| | Pteridospermatophyta | Gμ0_8 | 0.061300197 | 0.080139895 | -0.067706745 | 0.272991238 |
| | Sphenophyta | Gμ0_9 | 0.083477907 | 0.096753362 | -0.047873946 | 0.319054932 |
| | CO2 | Gμ0_10 | 0.000115653 | 0.000466773 | -0.000235566 | 0.002701865 |
| | O2 | Gμ0_12 | -2.896962776 | -3.08025556 | -8.434509064 | 1.055132805 |
| | Temp. | Gμ0_13 | -0.016744231 | -0.059261733 | -0.389656507 | 0.095364649 |
| | Total area covered by sed. | Gμ0_14 | -0.01431632 | -0.130011199 | -1.865529569 | 0.94933382 |
| Shrinkage weights (origination) | Diversity dependence | ωλ0_0 | 0.817399481 | 0.769017985 | 0.286618602 | 0.992297551 |
| | Plant diversity | ωλ0_1 | 0.630691792 | 0.551654442 | 0.002094385 | 0.98991706 |
| | Coniferophyta | ωλ0_2 | 0.427908472 | 0.44371059 | 0.001031803 | 0.978212323 |
| | Cycadophyta | ωλ0_3 | 0.285850682 | 0.364474434 | 0.00077293 | 0.97002265 |
| | Ginkgophyta | ωλ0_4 | 0.332319042 | 0.397784095 | 0.000774345 | 0.979012024 |
| | Gymnospermae | ωλ0_5 | 0.25674404 | 0.350559655 | 0.000514568 | 0.968381701 |

| | | | | | | |
|---|---|---|---|---|---|---|
| | Lycophyta | ωλ0_6 | 0.906748186 | 0.758804564 | 0.009861748 | 0.99748753 |
| | Pteridophyta | ωλ0_7 | 0.348923451 | 0.408344739 | 0.000157159 | 0.979469603 |
| | Pteridospermatophyta | ωλ0_8 | 0.39790116 | 0.434045071 | 0.000848299 | 0.981919219 |
| | Sphenophyta | ωλ0_9 | 0.428432589 | 0.446413763 | 0.001330723 | 0.983324222 |
| | CO2 | ωλ0_10 | 0.399066968 | 0.438705624 | 0.00054613 | 0.982272267 |
| | O2 | ωλ0_12 | 0.935565154 | 0.770409216 | 0.008893862 | 0.998157409 |
| | Temp. | ωλ0_13 | 0.955167485 | 0.766347028 | 0.007297152 | 0.999209434 |
| | Total area covered by sed. | ωλ0_14 | 0.50869759 | 0.50091278 | 0.00156799 | 0.991882751 |
| Shrinkage weights (extinction) | Diversity dependence | ωμ0_0 | 0.738668396 | 0.696416578 | 0.205773758 | 0.989869026 |
| | Plant diversity | ωμ0_1 | 0.40669051 | 0.44562767 | 0.001051734 | 0.986501642 |
| | Coniferophyta | ωμ0_2 | 0.780299594 | 0.668868512 | 0.00839271 | 0.994698982 |
| | Cycadophyta | ωμ0_3 | 0.506207412 | 0.48928456 | 0.00211929 | 0.98337551 |
| | Ginkgophyta | ωμ0_4 | 0.433919928 | 0.46037007 | 0.000749664 | 0.990518717 |
| | Gymnospermae | ωμ0_5 | 0.348340942 | 0.405415166 | 0.001027375 | 0.973163386 |
| | Lycophyta | ωμ0_6 | 0.462311845 | 0.466063059 | 0.000868194 | 0.987896078 |
| | Pteridophyta | ωμ0_7 | 0.471509005 | 0.472054664 | 0.00145384 | 0.985890018 |
| | Pteridospermatophyta | ωμ0_8 | 0.816324872 | 0.660691087 | 0.004306329 | 0.995644249 |
| | Sphenophyta | ωμ0_9 | 0.835306862 | 0.665080882 | 0.003909426 | 0.996851882 |
| | CO2 | ωμ0_10 | 0.644762468 | 0.568571239 | 0.001893475 | 0.997334879 |
| | O2 | ωμ0_12 | 0.822602002 | 0.704662999 | 0.011668885 | 0.994685625 |
| | Temp. | ωμ0_13 | 0.533740941 | 0.511443178 | 0.001750268 | 0.992760791 |
| | Total area covered by sed. | ωμ0_14 | 0.486041561 | 0.483450448 | 0.001361877 | 0.991622457 |

**Table S16.** Correlation parameters and Shrinkage weights of vertebrate in 0-30°N using MBD model. Baseline origination and extinction rates (λ0 and μ0) and correlation parameters (Gλ and Gμ). The mean correlation parameters values were divided into pink (negative) and blue (positive). Shrinkage weights greater than 0.5 show significant evidence for correlation and are marked in orange. The strong support significant for bolded.

| Parameters | | | Median | Mean | CI_lower | CI_upper |
|---|---|---|---|---|---|---|
| Baseline rates | origination | λ0 | 0.433276388 | 0.526739489 | 0.084178827 | 1.499424274 |
| | extinction | μ0 | 0.305957775 | 0.394215855 | 0.042419464 | 1.246305082 |
| Correlation parameters to origination | Diversity dependence | Gλ0_0 | -2.541725577 | -2.547569947 | -5.185138612 | -0.049396824 |
| | Plant diversity | Gλ0_1 | -0.002614085 | -0.012604886 | -0.104622425 | 0.035786747 |
| | Coniferophyta | Gλ0_2 | 0.058897021 | 0.074471656 | -0.005393329 | 0.232779132 |
| | Cycadophyta | Gλ0_3 | 0.013382128 | 0.048422685 | -0.103215907 | 0.34473181 |
| | Ginkgophyta | Gλ0_4 | -0.01238143 | -0.041539566 | -0.240938943 | 0.025920311 |
| | Gymnospermae | Gλ0_5 | -0.005247809 | -0.020662526 | -0.165327447 | 0.056516804 |
| | Lycophyta | Gλ0_6 | 0.001076922 | 0.010478707 | -0.034100136 | 0.131949809 |
| | Pteridophyta | Gλ0_7 | -0.003285098 | -0.01367389 | -0.104562719 | 0.030979528 |
| | Pteridospermatophyta | Gλ0_8 | 0.001556381 | 0.012140129 | -0.071862593 | 0.159579607 |
| | Sphenophyta | Gλ0_9 | 0.001521768 | 0.011179363 | -0.07148259 | 0.152296021 |
| | CO2 | Gλ0_10 | 1.73E-06 | 2.60E-05 | -0.000500305 | 0.000720751 |
| | O2 | Gλ0_12 | 0.37844483 | 2.991696054 | -1.41310976 | 19.96187928 |
| | Temp. | Gλ0_13 | -0.010542117 | -0.113938042 | -1.088431209 | 0.077895296 |
| | Total area covered by sed. | Gλ0_14 | -0.01233961 | -0.082455185 | -1.316661506 | 0.794686584 |
| Correlation parameters to extinction | Diversity dependence | Gμ0_0 | -2.496667582 | -2.440458752 | -5.303286492 | 0.089958893 |
| | Plant diversity | Gμ0_1 | 0.001129129 | 0.00445434 | -0.054438208 | 0.072414609 |
| | Coniferophyta | Gμ0_2 | -0.002505917 | -0.012881862 | -0.127876104 | 0.055711898 |
| | Cycadophyta | Gμ0_3 | 0.000533914 | 0.006930788 | -0.182996839 | 0.232789465 |
| | Ginkgophyta | Gμ0_4 | 0.000199622 | 0.002295018 | -0.106647031 | 0.127251219 |
| | Gymnospermae | Gμ0_5 | 0.038801076 | 0.056677382 | -0.035135313 | 0.23109869 |
| | Lycophyta | Gμ0_6 | -0.00456831 | -0.018901617 | -0.117636376 | 0.016197546 |
| | Pteridophyta | Gμ0_7 | 0.006331666 | 0.027772015 | -0.021381643 | 0.17648327 |
| | Pteridospermatophyta | Gμ0_8 | 0.00182277 | 0.012888656 | -0.072198551 | 0.155053902 |
| | Sphenophyta | Gμ0_9 | 0.002120835 | 0.013224749 | -0.071306099 | 0.149706495 |
| | CO2 | Gμ0_10 | 4.52E-08 | 1.81E-05 | -0.000553061 | 0.000685484 |
| | O2 | Gμ0_12 | -0.943078666 | -2.127328194 | -10.62609933 | 0.822343718 |
| | Temp. | Gμ0_13 | -0.002054564 | -0.011080196 | -0.2470847 | 0.190767111 |
| | Total area covered by sed. | Gμ0_14 | -0.007368436 | -0.08030723 | -1.361559687 | 0.846933073 |
| Shrinkage weights (origination) | Diversity dependence | ωλ0_0 | 0.844471917 | 0.74630269 | 0.038963969 | 0.995063084 |
| | Plant diversity | ωλ0_1 | 0.303714261 | 0.390305029 | 0.000428686 | 0.984910468 |
| | Coniferophyta | ωλ0_2 | 0.72289815 | 0.605697907 | 0.002539088 | 0.995174954 |
| | Cycadophyta | ωλ0_3 | 0.304064277 | 0.387465924 | 0.000549048 | 0.980728143 |
| | Ginkgophyta | ωλ0_4 | 0.402162274 | 0.45388164 | 0.000673139 | 0.992119264 |
| | Gymnospermae | ωλ0_5 | 0.234974879 | 0.339413856 | 0.000329743 | 0.968487889 |
| | Lycophyta | ωλ0_6 | 0.329921208 | 0.40933964 | 0.000484939 | 0.990659637 |

| | | | | | | |
|---|---|---|---|---|---|---|
| | Pteridophyta | ωλ0_7 | 0.344819106 | 0.411051957 | 0.000749488 | 0.982612615 |
| | Pteridospermatophyta | ωλ0_8 | 0.278740771 | 0.37785081 | 0.000457405 | 0.981692179 |
| | Sphenophyta | ωλ0_9 | 0.291602598 | 0.380984023 | 0.000420106 | 0.981604532 |
| | CO2 | ωλ0_10 | 0.286725018 | 0.385255647 | 0.0004965 | 0.982478991 |
| | O2 | ωλ0_12 | 0.439918779 | 0.482665794 | 0.000625854 | 0.997213187 |
| | Temp. | ωλ0_13 | 0.397956244 | 0.458217326 | 0.000839221 | 0.997747455 |
| | Total area covered by sed. | ωλ0_14 | 0.312515829 | 0.394740217 | 0.000557339 | 0.984826629 |
| | Diversity dependence | ωμ0_0 | 0.836619046 | 0.715714254 | 0.009412651 | 0.994469851 |
| | Plant diversity | ωμ0_1 | 0.294614284 | 0.380904132 | 0.000428083 | 0.977154817 |
| | Coniferophyta | ωμ0_2 | 0.277719273 | 0.375868374 | 0.000523715 | 0.977183999 |
| | Cycadophyta | ωμ0_3 | 0.250898153 | 0.353082198 | 0.000363172 | 0.970871891 |
| | Ginkgophyta | ωμ0_4 | 0.297410814 | 0.38741459 | 0.000472212 | 0.98324799 |
| | Gymnospermae | ωμ0_5 | 0.44553776 | 0.458405146 | 0.001244778 | 0.984223955 |
| Shrinkage weights (extinction) | Lycophyta | ωμ0_6 | 0.389298832 | 0.448776138 | 0.000491655 | 0.992537832 |
| | Pteridophyta | ωμ0_7 | 0.426807485 | 0.462321084 | 0.000391189 | 0.993580065 |
| | Pteridospermatophyta | ωμ0_8 | 0.296224606 | 0.386993925 | 0.000423576 | 0.978089616 |
| | Sphenophyta | ωμ0_9 | 0.290401492 | 0.384777312 | 0.000461285 | 0.981763304 |
| | CO2 | ωμ0_10 | 0.299068913 | 0.386499939 | 0.000455525 | 0.983774041 |
| | O2 | ωμ0_12 | 0.546933953 | 0.508828305 | 0.000832561 | 0.994608703 |
| | Temp. | ωμ0_13 | 0.308378165 | 0.401894712 | 0.000365342 | 0.986665298 |
| | Total area covered by sed. | ωμ0_14 | 0.299677667 | 0.393767369 | 0.000382899 | 0.986981781 |

**Table S17.** Correlation parameters and Shrinkage weights of vertebrate in 30-60°N using MBD model. Baseline origination and extinction rates (λ0 and μ0) and correlation parameters (Gλ and Gμ). The mean correlation parameters values were divided into pink (negative) and blue (positive). Shrinkage weights greater than 0.5 show significant evidence for correlation and are marked in orange. The strong support significant for bolded.

| Parameters | | | Median | Mean | CI_lower | CI_upper |
|---|---|---|---|---|---|---|
| Baseline rates | origination | λ0 | 0.36660914 | 0.437386123 | 0.077484819 | 1.18153427 |
| | extinction | μ0 | 0.183522889 | 0.280638184 | 0.02000237 | 1.185894073 |
| Correlation parameters to origination | Diversity dependence | Gλ0_0 | -4.215926617 | -4.297639404 | -7.239881745 | -1.958234054 |
| | Plant diversity | Gλ0_1 | 0.00022109 | 0.00141517 | -0.047149393 | 0.051476166 |
| | Coniferophyta | Gλ0_2 | 0.000220176 | 0.000729173 | -0.053270091 | 0.052028173 |
| | Cycadophyta | Gλ0_3 | 0.003956863 | 0.021237099 | -0.079073215 | 0.204163786 |
| | Ginkgophyta | Gλ0_4 | 0.001470417 | 0.00653444 | -0.040548778 | 0.073560601 |
| | Gymnospermae | Gλ0_5 | 0.006664249 | 0.02048884 | -0.048249256 | 0.142774439 |
| | Lycophyta | Gλ0_6 | 9.44E-05 | 0.001299421 | -0.022406217 | 0.028934013 |
| | Pteridophyta | Gλ0_7 | -7.44E-05 | -0.001339567 | -0.045262297 | 0.036029231 |
| | Pteridospermatophyta | Gλ0_8 | 0.003507273 | 0.017831853 | -0.045384577 | 0.132660258 |
| | Sphenophyta | Gλ0_9 | 0.004627001 | 0.016653076 | -0.045960399 | 0.118676147 |
| | CO2 | Gλ0_10 | -3.11E-06 | -5.40E-05 | -0.000903702 | 0.000374659 |
| | O2 | Gλ0_12 | 0.251453759 | 0.943394808 | -1.279482224 | 5.902329799 |
| | Temp. | Gλ0_13 | 0.000998849 | 0.009081856 | -0.106595604 | 0.163607031 |
| | Total area covered by sed. | Gλ0_14 | 0.002681946 | 0.029497491 | -0.578381319 | 0.722138324 |
| Correlation parameters to extinction | Diversity dependence | Gμ0_0 | -0.124977291 | -0.350017779 | -2.275855504 | 1.045321275 |
| | Plant diversity | Gμ0_1 | 0.012952117 | 0.039235056 | -0.020408224 | 0.199179742 |
| | Coniferophyta | Gμ0_2 | 0.000156456 | -0.004706125 | -0.13617393 | 0.075089114 |
| | Cycadophyta | Gμ0_3 | -0.000725722 | -0.007591692 | -0.163797637 | 0.11237045 |
| | Ginkgophyta | Gμ0_4 | 8.71E-05 | 0.001651537 | -0.052046507 | 0.065317465 |
| | Gymnospermae | Gμ0_5 | 0.010487041 | 0.03001793 | -0.062291489 | 0.179586118 |
| | Lycophyta | Gμ0_6 | -0.00061855 | -0.005285513 | -0.059509232 | 0.024222849 |
| | Pteridophyta | Gμ0_7 | 0.000969658 | 0.005903606 | -0.031585926 | 0.068506623 |
| | Pteridospermatophyta | Gμ0_8 | 0.005414958 | 0.038531438 | -0.042197179 | 0.303440351 |
| | Sphenophyta | Gμ0_9 | 0.0054768 | 0.042000615 | -0.043374961 | 0.310676708 |
| | CO2 | Gμ0_10 | -7.04E-06 | -0.000102224 | -0.001221158 | 0.000361695 |
| | O2 | Gμ0_12 | -0.679507301 | -2.885324599 | -14.15256496 | 0.876992817 |
| | Temp. | Gμ0_13 | -0.004688117 | -0.036582157 | -0.369530862 | 0.092632181 |
| | Total area covered by sed. | Gμ0_14 | -0.03367943 | -0.330967643 | -2.979096616 | 0.513680233 |
| Shrinkage weights (origination) | Diversity dependence | ωλ0_0 | 0.929831936 | 0.888175506 | 0.55581005 | 0.997244636 |
| | Plant diversity | ωλ0_1 | 0.174502842 | 0.294951068 | 0.000124615 | 0.949449744 |
| | Coniferophyta | ωλ0_2 | 0.135030932 | 0.251298975 | 2.77E-05 | 0.929387954 |
| | Cycadophyta | ωλ0_3 | 0.138136481 | 0.267053234 | 7.80E-05 | 0.932581357 |
| | Ginkgophyta | ωλ0_4 | 0.168866914 | 0.290104469 | 0.000119222 | 0.947885173 |
| | Gymnospermae | ωλ0_5 | 0.159109861 | 0.283028233 | 0.000120632 | 0.943656275 |
| | Lycophyta | ωλ0_6 | 0.137539171 | 0.259391847 | 6.86E-05 | 0.93555956 |

| | | | | | | |
|---|---|---|---|---|---|---|
| | Pteridophyta | ωλ0_7 | 0.142777766 | 0.275968877 | 8.47E-05 | 0.947139519 |
| | Pteridospermatophyta | ωλ0_8 | 0.229611114 | 0.33998862 | 0.000195333 | 0.972882712 |
| | Sphenophyta | ωλ0_9 | 0.209697632 | 0.331791772 | 0.000106639 | 0.969588261 |
| | CO2 | ωλ0_10 | 0.185643637 | 0.322561401 | 7.15E-05 | 0.976196903 |
| | O2 | ωλ0_12 | 0.253392137 | 0.372607245 | 0.000138619 | 0.979800884 |
| | Temp. | ωλ0_13 | 0.175596241 | 0.306861565 | 0.000111997 | 0.972176372 |
| | Total area covered by sed. | ωλ0_14 | 0.174908252 | 0.29780463 | 0.000115322 | 0.955889086 |
| Shrinkage weights (extinction) | Diversity dependence | ωμ0_0 | 0.219448762 | 0.316486288 | 0.000137852 | 0.944500125 |
| | Plant diversity | ωμ0_1 | 0.481631098 | 0.480601298 | 0.00025388 | 0.993349382 |
| | Coniferophyta | ωμ0_2 | 0.212268175 | 0.333662964 | 0.000196758 | 0.970735334 |
| | Cycadophyta | ωμ0_3 | 0.139749784 | 0.262086764 | 6.59E-05 | 0.938251474 |
| | Ginkgophyta | ωμ0_4 | 0.144384554 | 0.272452104 | 9.90E-05 | 0.945417859 |
| | Gymnospermae | ωμ0_5 | 0.244319351 | 0.342762569 | 0.000141106 | 0.961862853 |
| | Lycophyta | ωμ0_6 | 0.194494143 | 0.327116219 | 0.000116224 | 0.967386561 |
| | Pteridophyta | ωμ0_7 | 0.175527891 | 0.306260198 | 0.000137011 | 0.961179801 |
| | Pteridospermatophyta | ωμ0_8 | 0.269446918 | 0.388087372 | 0.000120751 | 0.991577703 |
| | Sphenophyta | ωμ0_9 | 0.269453522 | 0.397088189 | 0.000125025 | 0.991816678 |
| | CO2 | ωμ0_10 | 0.209435738 | 0.344684177 | 9.47E-05 | 0.984341979 |
| | O2 | ωμ0_12 | 0.427766677 | 0.477155349 | 0.000236133 | 0.996794049 |
| | Temp. | ωμ0_13 | 0.223391767 | 0.354541159 | 0.00011544 | 0.986468836 |
| | Total area covered by sed. | ωμ0_14 | 0.266301491 | 0.390938508 | 0.000105344 | 0.994391728 |

**Table S18.** Correlation parameters and Shrinkage weights of vertebrate in 30-60°S using MBD model. Baseline origination and extinction rates (λ0 and μ0) and correlation parameters (Gλ and Gμ). The mean correlation parameters values were divided into pink (negative) and blue (positive). Shrinkage weights greater than 0.5 show significant evidence for correlation and are marked in orange. The strong support significant for bolded.

| Parameters | | | Median | Mean | CI_lower | CI_upper |
|---|---|---|---|---|---|---|
| Baseline rates | origination | λ0 | 0.328510757 | 0.388975179 | 0.066559416 | 1.007580706 |
| | extinction | μ0 | 0.215869158 | 0.280458269 | 0.048301788 | 0.870352642 |
| Correlation parameters to origination | Diversity dependence | Gλ0_0 | -2.049698439 | -2.039445433 | -3.004063455 | -0.957456379 |
| | Plant diversity | Gλ0_1 | 0.003886565 | 0.010611989 | -0.011811687 | 0.060197377 |
| | Coniferophyta | Gλ0_2 | 0.000719958 | 0.003238456 | -0.049638335 | 0.056588436 |
| | Cycadophyta | Gλ0_3 | 0.0013143 | 0.005387844 | -0.071881924 | 0.099532881 |
| | Ginkgophyta | Gλ0_4 | 0.000365943 | 0.001318196 | -0.031841486 | 0.034215918 |
| | Gymnospermae | Gλ0_5 | 0.000145031 | -0.000251934 | -0.061243992 | 0.049226101 |
| | Lycophyta | Gλ0_6 | 0.001091154 | 0.005892224 | -0.009471875 | 0.051670504 |
| | Pteridophyta | Gλ0_7 | 0.001313731 | 0.005033796 | -0.014638363 | 0.04040064 |
| | Pteridospermatophyta | Gλ0_8 | 0.000991466 | 0.004417183 | -0.030181632 | 0.049640139 |
| | Sphenophyta | Gλ0_9 | 0.001031242 | 0.004209477 | -0.02859603 | 0.050338465 |
| | CO2 | Gλ0_10 | -6.90E-07 | -4.47E-06 | -0.000354096 | 0.000334376 |
| | O2 | Gλ0_12 | 0.114437337 | 0.427660634 | -0.593310954 | 3.10924726 |
| | Temp. | Gλ0_13 | 0.000220895 | -0.000591454 | -0.085613602 | 0.074506814 |
| | Total area covered by sed. | Gλ0_14 | 0.000434983 | -0.007734305 | -0.586751297 | 0.454042575 |
| Correlation parameters to extinction | Diversity dependence | Gμ0_0 | -0.302710469 | -0.428111183 | -1.562002796 | 0.201401216 |
| | Plant diversity | Gμ0_1 | -0.000209266 | -0.001419308 | -0.043019442 | 0.036045167 |
| | Coniferophyta | Gμ0_2 | -0.000207881 | -0.002092977 | -0.053774541 | 0.038481108 |
| | Cycadophyta | Gμ0_3 | -0.009186964 | -0.027334592 | -0.155760862 | 0.035678894 |
| | Ginkgophyta | Gμ0_4 | -0.001089504 | -0.007284541 | -0.068864288 | 0.023533668 |
| | Gymnospermae | Gμ0_5 | 0.085103847 | 0.084477476 | -0.007330446 | 0.207822905 |
| | Lycophyta | Gμ0_6 | -0.001058511 | -0.004159625 | -0.033743364 | 0.012735332 |
| | Pteridophyta | Gμ0_7 | -0.000458403 | -0.003009627 | -0.03642871 | 0.016669896 |
| | Pteridospermatophyta | Gμ0_8 | 0.005373603 | 0.022989747 | -0.02439929 | 0.143735035 |
| | Sphenophyta | Gμ0_9 | 0.007353799 | 0.028924399 | -0.023560996 | 0.163128789 |
| | CO2 | Gμ0_10 | 7.84E-07 | 2.34E-05 | -0.000304365 | 0.0004968 |
| | O2 | Gμ0_12 | -0.004040784 | -0.05009995 | -1.734061426 | 1.389804627 |
| | Temp. | Gμ0_13 | -0.004681676 | -0.025742195 | -0.194984677 | 0.036129269 |
| | Total area covered by sed. | Gμ0_14 | -0.01016632 | -0.075635526 | -0.818853978 | 0.302959339 |
| Shrinkage weights (origination) | Diversity dependence | ωλ0_0 | 0.771257775 | 0.727955845 | 0.246773589 | 0.989084859 |
| | Plant diversity | ωλ0_1 | 0.124217504 | 0.261236515 | 0.000131585 | 0.945319101 |
| | Coniferophyta | ωλ0_2 | 0.072180215 | 0.204064443 | 4.04E-05 | 0.905167119 |
| | Cycadophyta | ωλ0_3 | 0.06345826 | 0.174202173 | 5.20E-05 | 0.854792271 |
| | Ginkgophyta | ωλ0_4 | 0.06681807 | 0.18149359 | 7.66E-05 | 0.874159593 |
| | Gymnospermae | ωλ0_5 | 0.056754435 | 0.161068438 | 6.09E-05 | 0.841072319 |
| | Lycophyta | ωλ0_6 | 0.089701068 | 0.236799143 | 5.83E-05 | 0.950536934 |

| | | | | | | |
|---|---|---|---|---|---|---|
| | Pteridophyta | ωλ0_7 | 0.091596002 | 0.218368276 | 7.85E-05 | 0.920417533 |
| | Pteridospermatophyta | ωλ0_8 | 0.071210764 | 0.192145305 | 5.32E-05 | 0.888171114 |
| | Sphenophyta | ωλ0_9 | 0.066827521 | 0.188551464 | 9.29E-05 | 0.891220842 |
| | CO2 | ωλ0_10 | 0.085359643 | 0.223731063 | 0.000111877 | 0.933952654 |
| | O2 | ωλ0_12 | 0.081751237 | 0.219619328 | 6.62E-05 | 0.920694281 |
| | Temp. | ωλ0_13 | 0.075898412 | 0.200568092 | 4.00E-05 | 0.923771385 |
| | Total area covered by sed. | ωλ0_14 | 0.080075522 | 0.217231341 | 6.36E-05 | 0.935147701 |
| Shrinkage weights (extinction) | Diversity dependence | ωμ0_0 | 0.16467213 | 0.270999594 | 9.81E-05 | 0.921641459 |
| | Plant diversity | ωμ0_1 | 0.085024976 | 0.220819543 | 8.12E-05 | 0.928170678 |
| | Coniferophyta | ωμ0_2 | 0.076735609 | 0.201724124 | 0.000107385 | 0.905621544 |
| | Cycadophyta | ωμ0_3 | 0.084986132 | 0.213965664 | 5.23E-05 | 0.912124477 |
| | Ginkgophyta | ωμ0_4 | 0.085703655 | 0.217706244 | 5.15E-05 | 0.921021823 |
| | Gymnospermae | ωμ0_5 | 0.578359737 | 0.526314871 | 0.002220107 | 0.982007934 |
| | Lycophyta | ωμ0_6 | 0.095676208 | 0.230098621 | 7.50E-05 | 0.925155376 |
| | Pteridophyta | ωμ0_7 | 0.077960867 | 0.203956267 | 6.68E-05 | 0.914763632 |
| | Pteridospermatophyta | ωμ0_8 | 0.165571599 | 0.309094531 | 7.44E-05 | 0.971124016 |
| | Sphenophyta | ωμ0_9 | 0.183918722 | 0.334775375 | 9.21E-05 | 0.98018717 |
| | CO2 | ωμ0_10 | 0.090981102 | 0.236336464 | 9.10E-05 | 0.952622208 |
| | O2 | ωμ0_12 | 0.065160286 | 0.189425277 | 8.21E-05 | 0.889555274 |
| | Temp. | ωμ0_13 | 0.112162482 | 0.26932821 | 9.00E-05 | 0.963030009 |
| | Total area covered by sed. | ωμ0_14 | 0.093375396 | 0.233720959 | 6.68E-05 | 0.948114548 |

**Table S19.** Genus-level extinction rates and diversity changes of different clades across the Permian–Triassic boundary.

| Genus | | Extinction rates cross PTB (genus) | Diversity loss | Diversity loss from | Diversity loss to | Duration (Ma) | References |
|---|---|---|---|---|---|---|---|
| Global macroplants | | 0.2424 | 6.70% | 254 | 251 | 3 | This study |
| Global sporomorphs | | 0.225 | 8.60% | 251.6 | 249.8 | 1.8 | This study |
| Global vertebrates | | 1.1022 | 66.70% | 257 | 251 | 6 | This study |
| Global vertebrates | | | ~52.68% | Late Changhsingian-Induan | | | Benton and Newell, 2014 |
| Global insects | | 0.57 | 62.69% | 252 | 250 | 2 | Jouault et al., 2022 |
| Global marine organisms | | | ~72% | 252.73 | 251.95 | 0.78 | Fan et al., 2020 |
| Global marine organisms | | 1.73 | | | | | Foote, 2003 |
| Global marine organisms | | 2.12 | | | | | Kocsis et al., 2019 |
| Global marine organisms | | 1.5 | | | | | Alroy, 2008 |
| Plant latitudes | 0-30°N | 0.0943 | 33.30% | 255.6 | 250.6 | 5 | This study |
| | 30-60°N | 0.1765 | | | | | This study |
| | 60-90°N | 0.4675 | 87.50% | 253.9 | 249.3 | 4.6 | This study |
| | 0-30°S | 0.5352 | 84.38% | 252.12 | 249.7 | 2.42 | This study |
| | 30-60°S | 0.0607 | | | | | This study |
| | 60-90°S | 0.1619 | | | | | This study |
| Vertebrate latitudes | 0-30°N | 0.3515 | | | | | This study |
| | 30-60°N | 0.6534 | 75.00% | 255.5 | 253.2 | 2.3 | This study |
| | 30-60°S | 0.4781 | 60% | 253.7 | 251.9 | 1.8 | This study |
| Phytogeographical Zones | Angara | 0.1633 | 76.20% | 249 | 240 | 9 | This study |
| | Euramerica | 0.0115 | 16.70% | 251.8 | 245.5 | 6.3 | This study |
| | Gondwana | 0.1438 | | | | | This study |

| | | | | | | | |
|---|---|---|---|---|---|---|---|
| | South China | 0.3833 | 50.82% | 254.1 | 249 | 5.1 | This study |

**Table S20.** Species-level extinction rates and diversity changes of different clades across the Permian–Triassic boundary.

| Species | Extinction rates cross PTB (species) | Diversity loss | Diversity loss from | Diversity loss to | Duration (Ma) | References |
|---|---|---|---|---|---|---|
| Global macroplants | 0.56 | 19.80% | 252 | 249 | 7 | This study |
| Global macroplants | | 20.08% | | | | Niklas et al., 1985 |
| Global vertebrates | 1.1006 | 66.18% | 253.1 | 251.4 | 1.7 | This study |
| Global marine organisms | | ~83.3% | 252.73 | 251.95 | 0.78 | Fan et al., 2020 |
